# Early Pregnancy Marks Significant Shifts in the Oral Microbiome

**DOI:** 10.1101/2025.09.29.679276

**Authors:** Shani Finkelstein, Sigal Frishman, Sondra Turjeman, Oshrit Shtossel, Evgeny Tikhonov, Meital Nuriel-Ohayon, Yishay Pinto, Polina Popova, Alexandra S. Tkachuk, Elena A. Vasukova, Anna D. Anopova, Evgenii A. Pustozerov, Tatiana M. Pervunina, Elena N. Grineva, Moshe Hod, Betty Schwartz, Eran Hadar, Omry Koren, Yoram Louzoun

## Abstract

**Background:** Most studies of the oral microbiome during pregnancy have focused on the second and third trimesters (T2, T3, respectively). Findings remain inconsistent-some report shifts in specific taxa, whereas others observe little change in diversity. To date, no large-scale longitudinal study has examined oral microbiome development across all three trimesters, leaving early gestational dynamics (during the first trimester; T1) largely unexplored.

**Methods:** We conducted a longitudinal analysis of the oral microbiome in 346 pregnant women from Israel and validated key findings in an independent cohort of 154 pregnant women from Russia. In Israel, saliva samples were collected during T1 (11-14 weeks), T2 (24-28 weeks), and T3 (32-38 weeks) trimesters; in Russia, samples were collected during T2 and T3 with similar ranges of gestational weeks. Alongside sample collection, participants completed dietary and health questionnaires to assess maternal factors that could influence microbial composition. Microbial profiles were analyzed to test for (i) differential abundance across trimesters and (ii) the influence of maternal nutrition and lifestyle factors on these dynamics.

**Results:** Significant shifts in oral microbial composition were observed as early as the transition from T1 to T2. Alpha diversity decreased progressively across pregnancy (Shannon index: T1 = 3.261, T2 = 3.173, T3 = 3.109; Kruskal-Wallis p = 0.0023). Notable taxonomic changes included a significant reduction in Verrucomicrobiota (particularly *Akkermansia muciniphila*) and an increase in Synergistota from T1 to T2 (adjusted *p* < 0.01), alongside an increase in Gammaproteobacteria and a decrease in Erysipelotrichia, suggesting an ecological shift towards potentially pro-inflammatory communities. Despite these systematic population-level changes, within-subject microbial distances across trimesters were smaller than between-subject distances, indicating that individual women maintained relatively stable microbial profiles over time. Among 54 maternal variables examined, gluten-free diet showed the strongest and most consistent associations with oral microbiome composition across all trimesters, followed by smoking history and conception method. Key findings were validated in an independent cohort of 154 Russian women.

**Conclusions:** This study provides the first large-scale evidence of significant oral microbiome changes beginning in early pregnancy, characterized by reduced diversity and a directional shift toward potentially pro-inflammatory communities. The strong associations with gluten consumption and smoking suggest a large-scale effect of lifestyle on the pregnancy oral microbiome. The alteration in the microbial composition highlights the oral microbiome as a sensitive marker of gestational physiology.

## 1 Introduction

Several reports have demonstrated that the oral microbiome undergoes distinct compositional changes throughout pregnancy, influenced by hormonal, immunologic, and metabolic shifts[7, 50, 6]. However, findings remain inconsistent across studies. While some studies report relatively stable overall microbial diversity, others observe a trend towards decreased microbial richness and shifts in community structure, particularly in the second (T2) and third (T3) trimesters[47, 7, 18].

Studies using 16S rRNA gene sequencing have shown significant increases in certain taxa in T2 compared to T3, such as the phylum Bacteroidota, and the genera *Veillonella* (notably *Veillonella dispar* and *Veillonella atypica*), and *Granulicatella*, with a concurrent decrease in *Streptococcus sanguinis* and *Selenomonas*[7, 10, 47]. T3 is associated with a further reduction in microbial richness and increased dominance of specific genera, including Bacteroidota (notably Prevotella spp.), Veillonella spp. (especially Veillonella dispar and Veillonella atypica), and Streptococcus mutans[7], suggesting a shift toward a more distinct, potentially pro-inflammatory community [42, 21]. These changes occur in parallel with an increase in sex hormone levels[33]. The oral microbiome tends to revert to a pre-pregnancy state within weeks postpartum[18]. Importantly, while the oral microbiome is more stable than the vaginal or gut microbiome [44], pregnancy-associated changes can increase susceptibility to oral disease and have been associated with adverse pregnancy outcomes[8]. Although the evolution of the oral microbiome in T2 and T3 has been extensively studied, to our knowledge, there are currently no large-scale studies based on next generation sequencing that look at the evolution of the oral microbiome from the first trimester (T1) of pregnancy onwards, nor any clear evidence of a change in the oral microbiome during this period (see table 2 in [18] and [3, 15]).

Diet is a central determinant of oral microbial ecology, shaping both community structure and function [35, 22]. Frequent intake of fermentable carbohydrates, particularly sugars, enriches acidogenic and aciduric taxa such as *Streptococcus mutans* and *Lactobacillus*, promoting dysbiosis and cariogenesis [27, 41]. Conversely, diets rich in fiber, plant polyphenols, and micronutrients, such as vitamin D and calcium, are associated with greater oral microbial diversity, resilience, and anti-inflammatory properties [51, 17]. In addition, nutritional deficiencies, including iron and folate, can also alter microbial colonization [4]. Thus, dietary patterns interact with host physiology to create either a protective or disease-prone oral environment.

During pregnancy, nutritional demands increase to support maternal health and fetal development, and dietary habits often change due to nausea, cravings, or medical recommendations. These shifts can exacerbate pregnancy-related changes in the oral microbiome. For instance, increased snacking and carbohydrate intake during the first trimester (T1) may favor cariogenic bacteria, while micronutrient supplementation (e.g., folic acid, iron) may mitigate dysbiosis [45, 28]. Restrictive diets such as gluten-free regimens, although necessary for individuals with celiac disease, can reduce dietary fiber and prebiotic intake, potentially leading to decreased microbial diversity and enrichment of opportunistic taxa in the oral cavity [29]. Together, these findings suggest that nutrition is a key, though underexplored, modifier of pregnancy-associated oral microbiome changes. Here, we analyzed the oral microbiome at the end of T1, T2, and T3 in 346 Israeli women [31], aiming to characterize how microbial composition and dynamics change across pregnancy stages in relation to maternal demographics, lifestyle, and dietary patterns. To assess the reproducibility of these findings, we validated our results in an independent cohort of 154 Russian women sampled during T2 and T3.

## 2 Methods

### 2.1 Study Population

To analyze the development of the oral microbiome during pregnancy, we recruited an Israeli cohort of pregnant women sampled during T1, T2, and T3. A total of 346 Israeli women were recruited, and 467 oral microbiome samples were processed: 235 from T1 (11-14 gestational weeks), 144 from T2 (24-28 gestational weeks), and 88 from T3 (32-38 gestational weeks; Table 1). It is important to note that a subset of participants was diagnosed with gestational diabetes mellitus in T2 (GDM; 79 cases and 267 controls). Full cohort details are provided in Pinto et al [31].

**Table 1:**
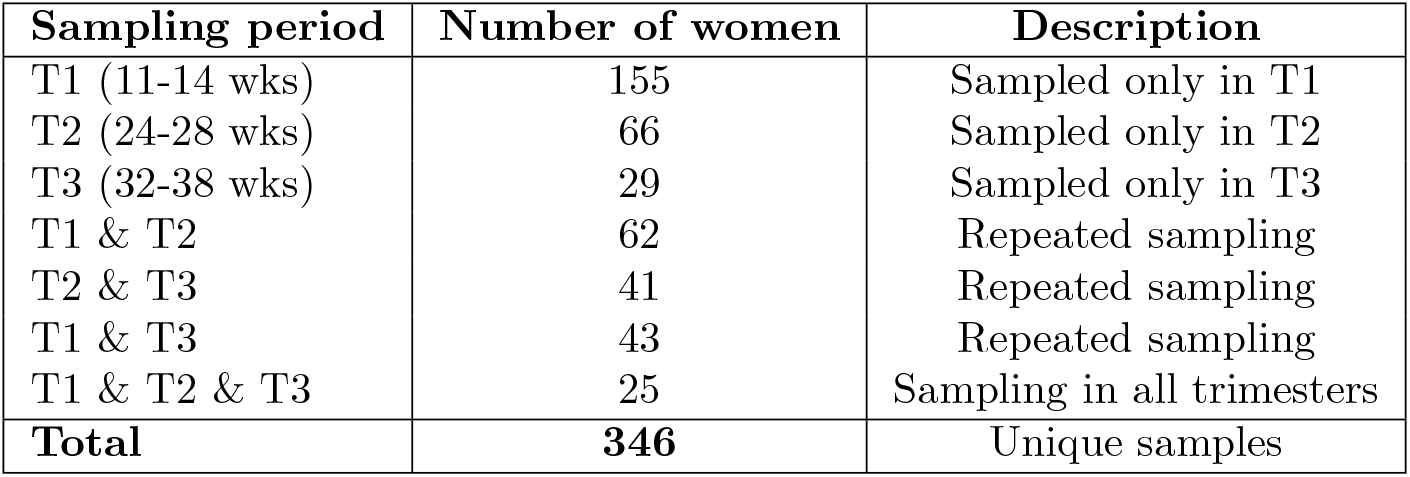
Distribution of Israeli cohort participants across trimesters and their overlap.

| Sampling period | Number of women | Description |
| --- | --- | --- |
| T1 (11-14 wks) | 155 | Sampled only in T1 |
| T2 (24-28 wks) | 66 | Sampled only in T2 |
| T3 (32-38 wks) | 29 | Sampled only in T3 |
| T1 & T2 | 62 | Repeated sampling |
| T2 & T3 | 41 | Repeated sampling |
| T1 & T3 | 43 | Repeated sampling |
| T1 & T2 & T3 | 25 | Sampling in all trimesters |
| **Total** | **346** | Unique samples |

A second cohort of pregnant women, a Russian cohort, was recruited to validate our findings; In total, 154 pregnant women were recruited, of whom 138 were sampled in T2 (16-28 gestational weeks) and 106 in T3 (34-37 gestational weeks), with a total of 244 oral microbiome samples.

The majority of participants were diagnosed with GDM (139 cases and 15 controls); full cohort details are provided in Popova et al. 2025[32].

### 2.2 Sample Collection

#### 2.2.1 Israeli cohort

At recruitment, the height and current weight of the study participants were recorded, and participants were interviewed by a dietitian for stress level, working hours, sleeping hours, smoking, education, and a 24-hour recall for food intake. Participants then provided a saliva sample, according to Human Microbiome Project protocols[1], collected in 1.5 ml tubes and stored at −80°C until processing. Similar procedures were repeated in the study population in T2 (24-28 gestational weeks) and T3 (32-38 gestational weeks) of pregnancy.

#### 2.2.2 Russian cohort

At recruitment (T2; 16-28 weeks of gestation), the participants’ pre-pregnancy height and weight were recorded, and participants were asked to fill out questionnaires regarding smoking habits and nutrition[32]. Participants then provided saliva, as described above. Samples were collected again in T3 (34-37 gestational weeks). It is important to note that the questionnaires used in this cohort differed from those administered in the Israeli cohort, and thus, for validation analyses, only the variables common to both cohorts were used.

### 2.3 Sample processing

DNA was extracted from all collected samples using the PowerSoil DNA Isolation Kit (MO BIO, Carlsbad, CA, USA) according to the manufacturer’s instructions and following a 2~min bead beating step (BioSpec, Bartlesville, OK, USA). The variable V4 region of the 16S rRNA gene was PCR-amplified using the 515F and 806R barcoded primers following the Earth Microbiome Project protocol[11]. Each PCR reaction contained 25 *µ*l with ~ 40 ng/*µ*l of DNA, 2 *µ*l 515F (forward, 10 *µ*M) primer, 2 *µ*l 806R (reverse, 10 *µ*M) primer, and 25 *µ*l PrimeSTAR Max PCR Readymix (Takara, Mountain View, CA, USA). PCR conditions were as follows: 30 cycles of denaturation at 98^°^C for 10 s, annealing at 55^°^C for 5 s, and extension at 72^°^C for 20 s, followed by a final elongation at 72^°^C for 1 min. Amplicons were purified using AMPure magnetic beads (Beckman Coulter, Indianapolis, IN, USA) and quantified using the Picogreen dsDNA quantitation kit (ThermoFisher, Waltham, MA, USA). Equimolar amounts of DNA from individual samples were pooled and sequenced using the Illumina MiSeq platform at the Azrieli Faculty of Medicine’s Genome Center.

### 2.4 Computational analysis and statistics

Sequencing data was run through the YaMAS pipeline (https://github.com/louzounlab/YaMAS)[38], which uses QIIME2 [9] for quality control and sequence processing. Quality control measures include filtering out low-quality reads, removing samples with fewer than 2,000 total reads, and trimming sequences. The filtered sequences were then processed to identify Amplicon Sequence Variants (ASVs), which were used to generate a table summarizing their abundance. Taxonomy was assigned with a Naive Bayes classifier trained on the GreenGenes 13 8 99% ASVs reference database[13]. The resulting ASV tables for the Israeli and Russian cohorts were then preprocessed separately using the MIPMLP pipeline[19], consisting of four steps: (1) merging similar features based on taxonomic classifications, (2) scaling the distribution, (3) standardizing features to z-scores, and (4) applying dimensionality reduction. Taxa were collapsed at the most resolved taxonomic level using the mean. All analyses were conducted in Python (v3.9). Analyses were performed separately for the Israeli and Russian cohorts to ensure independent validation.

Alpha diversity metrics (Shannon diversity index and observed richness) were computed from relative abundance tables using scikit-bio. Global differences in alpha diversity across trimesters were assessed using the Kruskal-Wallis test, followed by pairwise two-tailed Mann-Whitney U tests with FDR correction (T1 vs. T2, T2 vs. T3, and T1 vs. T3). Since the Shannon index inherently includes a logarithmic component, all computations were performed on relative (non-log-transformed) abundance data. To quantify within- and between-subject variability in microbial composition over time, we computed multiple distance metrics on abundance profiles: cosine distance, Bray-Curtis dissimilarity, Jaccard dissimilarity, and Euclidean distance. For each metric, abundance profiles were first log-transformed (to reduce the disproportionate influence of highly abundant taxa) and then z-score normalized (to standardize scale across samples).

For each subject with samples from multiple trimesters, pairwise distances were calculated between all trimester combinations with available data (T1-T2, T2-T3, T1-T3) using each metric. We compared three types of distance distributions: (1) intra-individual consecutive (same woman across consecutive trimesters: T1-T2 or T2-T3), (2) intra-individual non-consecutive (same woman across T1-T3), and (3) inter-individual (different women, within or across trimesters). Differences between distributions were evaluated using one-sided Mann-Whitney U tests to determine whether samples from the same woman were more similar to each other than samples from different women, indicating subject-specific temporal stability.

To assess overall differences in microbial community structure, we applied PERMANOVA (permutational multivariate analysis of variance, 9,999 permutations) as implemented in scikit-bio [34]. For the GDM analysis, two-way PERMANOVA was tested: (1) the effect of GDM status (GDM vs. Control), (2) the effect of trimester (T1, T2, T3), and (3) the Group × Time interaction. To determine which specific trimesters differed in microbial composition, post-hoc pairwise PERMANOVA comparisons were conducted between all timepoint pairs (T1 vs T2, T1 vs T3, T2 vs T3), with p-values adjusted for multiple testing using the Benjamini-Hochberg false discovery rate (FDR) correction. Stratified analyses by trimester were also performed to assess GDM effects at individual timepoints. Effect sizes were reported as *η*^2^ (proportion of variance explained)

To identify compositional changes at the phylum level, relative abundances were log-transformed to reduce the influence of highly abundant taxa, and differences across trimesters were assessed using two-sided Mann-Whitney U tests with FDR correction. For finer taxonomic resolution, two complementary frameworks were used: (1) GIMIC method [37], which identifies groups of taxa exhibiting coordinated changes between consecutive trimesters (T1 vs. T2 and T2 vs. T3), and (2) miMic test [36], which detects taxa with significant differences between trimesters (T1 vs. T2, T1 vs. T3, and T2 vs. T3) using Mann-Whitney U tests followed by FDR correction. This hierarchical approach allowed consistent evaluation of both collapsed (at class or phylum levels) and specific (species-level) changes over time.

Associations between microbial taxa and maternal metadata were analyzed within each trimester (T1, T2, T3) and longitudinally between consecutive trimesters (T1-T2, T2-T3). For longitudinal analyses, we computed the difference in microbial abundance between consecutive trimesters for each woman and correlated these differences with baseline metadata values.

Metadata included nutritional, demographic, and clinical features collected via standardized questionnaires (Table S1). Categorical variables were one-hot encoded to generate binary indicator variables. To avoid spurious correlations, we retained only features (including binary encoded categories) present in at least five participants and excluded features with no variance across the cohort. Continuous numerical variables were retained without additional filtering. For association testing, microbial relative abundances were log-transformed and z-score standardized (centered-log-ratio transformation) to ensure comparability across taxa. Spearman’s rank correlation was used for continuous metadata features, and point-biserial correlation for binary features. Multiple testing correction was performed separately for each maternal feature across all taxa using the FDR method (Benjamini-Hochberg), with significance determined at adjusted *p* < 0.05.

To assess whether maternal features with similar microbiome associations also correlate with each other, we performed an additional correlation analysis. For each maternal feature from the taxon-feature association analysis described above, we extracted its vector of correlation coefficients across all microbial taxa. We then computed Pearson correlations between these microbial correlation profiles across all pairs of maternal features, creating a feature-feature similarity matrix. This quantified whether features showing similar patterns of association with the microbiome also tend to co-vary in the metadata. Statistical significance was determined at *p* < 0.05 after FDR correction (Benjamini-Hochberg method). To validate the reproducibility of our findings, all analyses, including diversity metrics, PERMANOVA tests, GIMIC and miMic analyses, distance-based intra-individual variability, and taxon-feature correlations, were independently repeated in the Russian cohort using identical preprocessing, normalization, and statistical parameters. Because first-trimester (T1) samples were unavailable, analyses were limited to T2 and T3.

## 3 Results

We analyzed 467 saliva samples collected from 346 Israeli women across all three trimesters of pregnancy (Table 1, Figure 1A). In addition to 16S rRNA gene sequencing, each participant completed dietary and health questionnaires and underwent clinical tests (Table S1 for questionnaire details; Fig. S1 for feature distributions).

**Figure 1.**
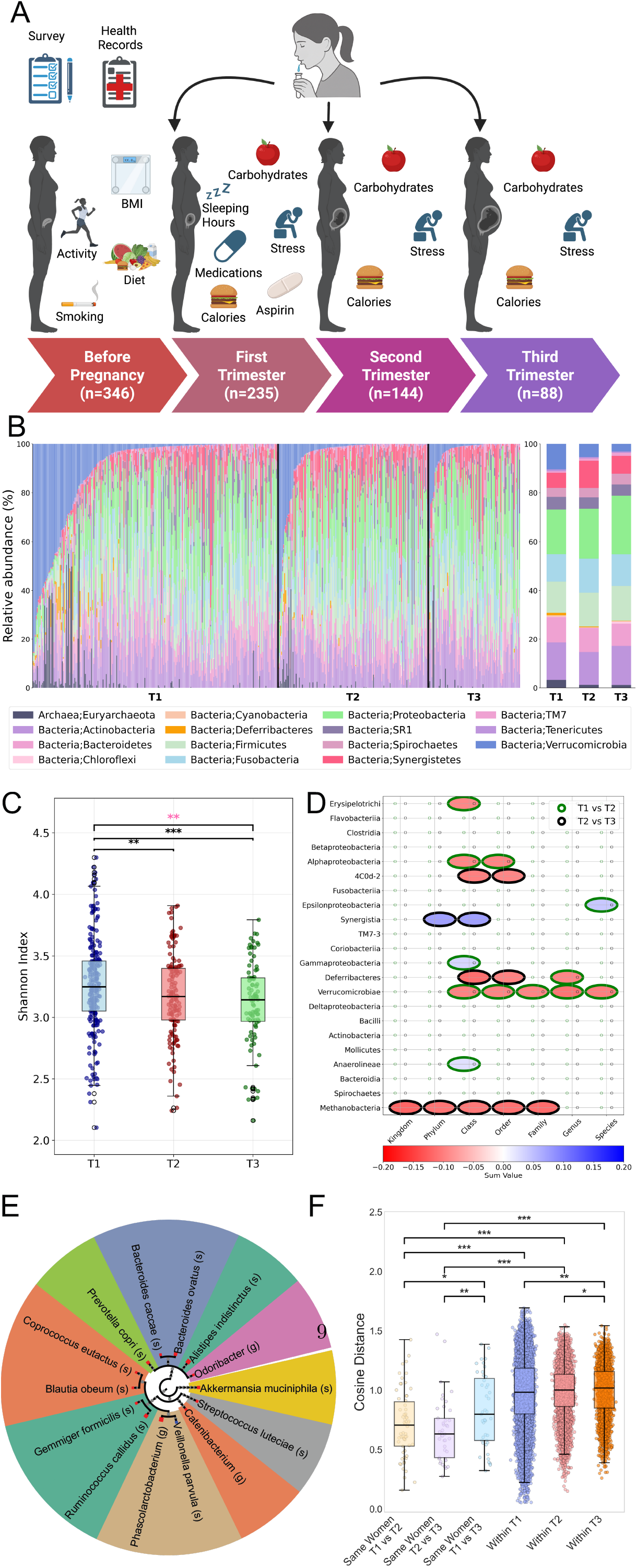
Longitudinal changes in the oral microbiome between the trimesters. (A) Schematic overview of the study design and sampling strategy for the oral microbiome analysis of 346 pregnant women across trimesters (T1, T2, T3). (B) Stacked bar plots of microbial composition at the phylum level, with each color representing a different phylum and each bar corresponding to one sample (summing to 100% relative abundance per sample). The average distribution on the right highlights a decrease in Verrucomicrobiota during pregnancy, particularly between T1 and T2 (*p* = 0.0011; see Results). (C) Alpha diversity (Shannon diversity index) boxplots, with dots representing individual samples, showing medians and interquartile ranges. Statistical significance is indicated by asterisks (\*\**p* < 0.01, Mann-Whitney test, T1 > T2; \*\*\**p* < 0.001, T1 > T3). A Kruskal-Wallis test confirmed overall group differences (*p* < 0.01, highlighted in pink). (D) GIMIC results depicting taxonomic shifts, with edge colors indicating comparisons: green for T1 vs. T2, black for T2 vs. T3. Bubbles summarize total differences across trimesters at each taxonomic level, with colors indicating enrichment (T1 vs. T2: red for T1, blue for T2; T2 vs. T3: red for T2, blue for T3), highlighting consistent shifts across taxonomic levels. (E) miMic test results identifying microbes with significant differences between T1 and T2, mapped onto a taxonomic tree. Dot colors indicate directionality (blue for increase in T2, red for decrease in T2), and dot size reflects the absolute Mann-Whitney test statistic multiplied by 30. Background shading groups taxa by family. (F) Cosine distance boxplots comparing microbiome dissimilarity across scenarios, with colors distinguishing comparison types: same women across trimesters, different women within trimesters, and different women between trimesters. Higher values indicate greater dissimilarity, with significance marked by asterisks (\**p* < 0.05, \*\**p* < 0.01, \*\*\**p* < 0.001). Distances between samples from the same woman were consistently smaller than in other scenarios, with reduced variance in distances between different women in T2 and T3. See Fig. S3 for miMic results for T1 vs. T3; no results were found for T2 vs. T3, and Fig. S4 for additional distance metrics.

We observed significant differences in community composition across pregnancy trimesters (PERMANOVA on Bray-Curtis distances: pseudo-F = 2.2, *p* = 0.001, *η*^2^ = 0.9%; Jaccard: pseudo-F = 4.4, *p* < 0.001, *η*^2^ = 1.6%; Fig. S2, S3), indicating substantial temporal shifts in the oral microbiome during gestation. Post-hoc pairwise comparisons with FDR correction revealed that the first trimester differed significantly from both the second (adjusted *p* = 0.006) and third trimesters (adjusted *p* = 0.003), whereas the second and third trimesters were compositionally similar (adjusted *p* = 0.052).

To ensure that these temporal patterns were not confounded by GDM status, we performed a two-way PERMANOVA testing for GDM effects and Group × Time interactions. GDM status explained minimal varaicne compared to trimester (Bray-Curtis: *η*^2^ = 0.3%, *p* = 0.094; Jaccard: *η*^2^ = 0.5%, *p* = 0.016). While Group × Time interactions were detected in both metrics (*p* ≤ 0.002), stratified analyses revealed metric-dependent patterns: Bray-Curtis showed a modest T2-specific difference (*p* = 0.041), whereas Jaccard showed no individual time point reached significance (all *p* > 0.25). Importantly, both groups exhibited qualitatively similar temporal progressions. Therefore, subsequent analyses focused on characterizing temporal dynamics across pregnancy.

To identify the taxa contributing to temporal community shifts, we first examined changes at the phylum level. We observed a significant increase in Synergistota from T1 to T2 and a significant decrease in Verrucomicrobiota over the same interval (adjusted *p* = 0.003 and *p* = 0.006, respectively) (Fig. 1B; Table S2 for mean and median phylum-level abundances).

Alpha diversity for all Israeli women, as quantified by the Shannon diversity index, exhibited significant differences across pregnancy trimesters (Kruskal-Wallis test, *p* = 0.0023). Pairwise Mann-Whitney tests indicated a significant decrease from T1 to T2 (adjusted *p* = 0.031) and from T1 to T3 (adjusted *p* = 0.010; Fig. 1C). Similar patterns were observed for species richness (Fig. S6). Shannon diversity did not differ significantly between T2 and T3 (adjusted *p* = 0.264); median values were 3.261 (T1), 3.173 (T2), and 3.109 (T3) (Fig. 1C).

To further characterize the microbial shifts driving these community-level changes, we examined differential taxonomic composition across trimesters and across taxonomic levels using GIMIC [37] (a method to represent differences at different taxonomic levels). Several taxa within the classes Alphaproteobacteria, Verrucomicrobiota, and Erysipelotrichia consistently decreased from T1 to T2, while Methanobacteria and Deferribacteres decreased from T2 to T3. The direction and magnitude of these shifts were stable across taxonomic levels, with Deferribacteres showing a consistent decline throughout pregnancy (Fig. 1D). To statistically test the difference, we used miMic (the statistical test matching GIMIC)[36] to detect specific taxa differing between trimesters (T1 vs. T2 and T1 vs. T3; Fig. 1E and Fig. S7). *A. muciniphila*, belonging to the Verrucomicrobiota phylum, which consistently decreased from T1 to T2 in the GIMIC analysis, also showed a significant species-level decrease (Fig. 1E). Other notable shifts included an increase in Gammaproteobacteria and a decrease in Erysipelotrichia in T2. No significant associations were observed between T2 and T3, indicating that the majority of detectable taxonmoic shifts occurred during early pregnancy (T1 to T2).

To assess the variability in microbial composition over time within and between women, we calculated cosine distances between samples. The distances between microbiomes of the same women across consecutive trimesters were significantly lower than those between different women, indicating subject-specific temporal stability in community composition (adjusted *p* < 0.001). However, when comparing non-consecutive trimesters, comparisons between the same women from T1 and T3 revealed significantly greater dissimilarity than T1-T2 or T2-T3 alone, consistent with cumulative compositional divergence that accumulates progressively over the course of pregnancy (T1-T2 vs. T1-T3: one-sided Mann-Whitney U = 1052, adjusted *p* = 0.033; T2-T3 vs. T1-T3: one-sided Mann-Whitney U = 596, adjusted *p* = 0.005; Fig. 1F). Similar patterns were observed for alternative distance metrics (Fig. S8).

We examined maternal factors associated with the oral microbiome development. The correlation between normalized microbial frequencies and 54 maternal variables collected pre-pregnancy or during specific trimesters, encompassing dietary habits (n=12), demographic factors (n=8), medical history (n=15), clinical measurements (n=14), and pregnancy outcomes (n=8) was computed (see Fig. 1A and Table S1 for complete feature descriptions).

Of these 54 features, three variables demonstrated consistent and significant correlations with the oral microbiome across trimesters after FDR correction, indicating that the majority of maternal features had little association with the oral community composition (Fig. 2A). The most robust associations were observed for adherence to a gluten-free diet, which correlated with oral microbiome structure in all three trimesters as well as with inter-trimester changes (T1-T2 and T2-T3). Tree-based association mapping (Fig. 2B) revealed that a gluten-free diet on T2 was linked to multiple taxa, with the strongest effects involving members of the families Lachnospiraceae and Ruminococcaceae.

**Figure 2.**
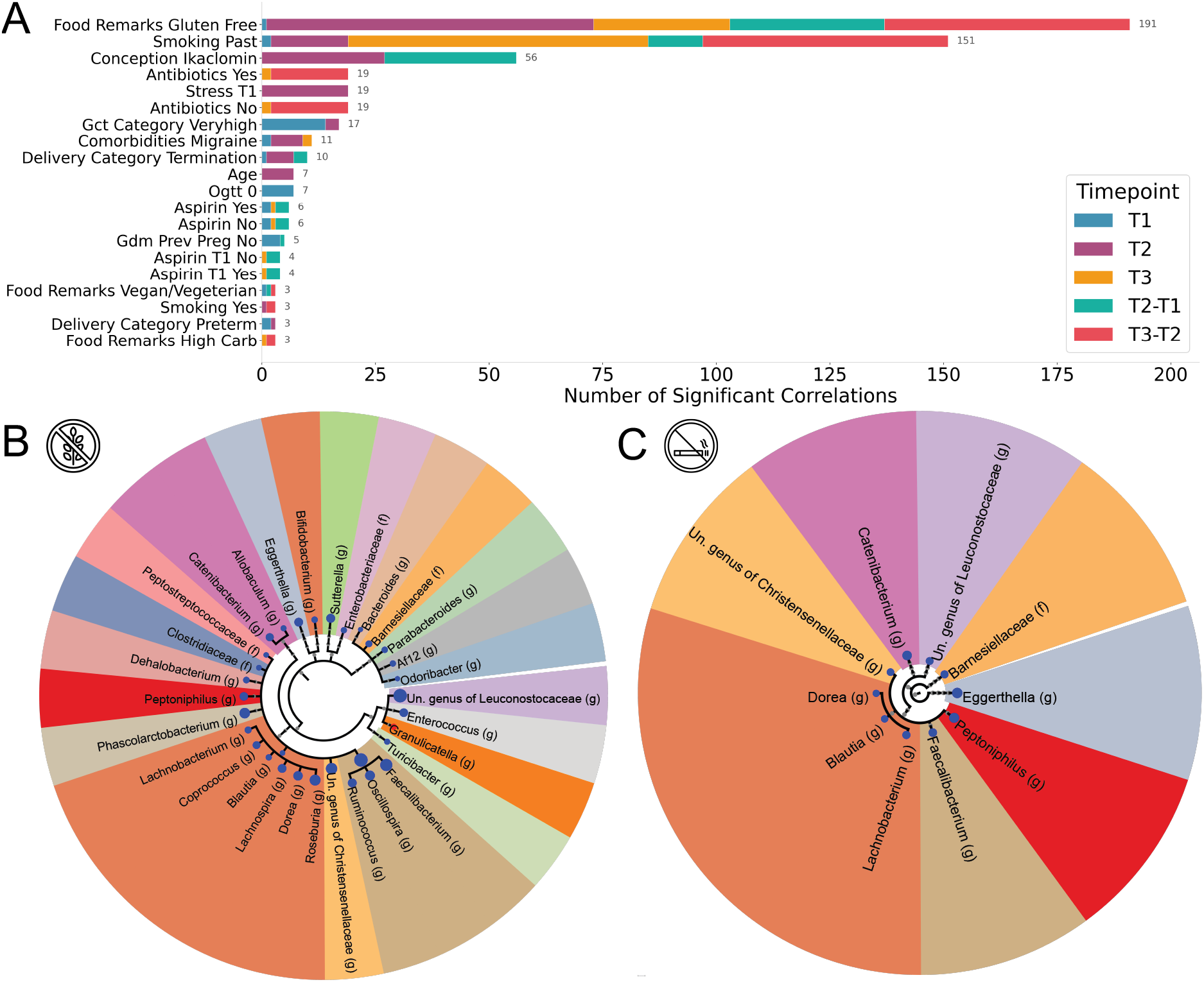
Associations between maternal characteristics and the oral microbiome. (A) Bar plot showing the number of significant correlations (FDR-corrected, *p* < 0.05) between maternal features and microbial taxa in T1 (blue), T2 (purple), and T3 (orange), as well as within-woman differences between consecutive trimesters (T2-T1, light blue; T3-T2, red). Only the top 20 features (ranked by total number of significant associations across all comparisons) are shown. (B)(C) Circular taxonomic tree mapping microbes significantly associated with gluten-free diet, and smoking history, respectively, in oral microbiome T2. Following the same format as the miMic tree in Fig. 1E, Dot colors indicate directionality (blue for increased abundance, red for decreased abundance in women following a gluten-free diet, smoked in the past), dot size reflects the absolute Mann-Whitney test statistic multiplied by 30, and background shading groups taxa by family. (B)(C) share the same family colors. The trees were filtered to show only the significant genera and species (See Fig. S9-S18 for additional full circular taxonomic trees for other features and trimesters, formatted similarly to the miMic tree).

A history of smoking was also significantly associated with oral microbial composition. Most associations were in taxa within Clostridiales and Bacteroidales classes (Fig. 2A; with T2 associations shown in Fig. 2C). These associations were present in all trimesters and with intertrimester changes, suggesting that smoking history exerts a lasting effect on the maternal oral microbiome.

Ikaclomin Conception method was likewise associated with oral microbial composition, sharing associations with taxa from the orders Clostridiales and Bacteroidales to Actinomycetales, in T2 (Fig. 2A, Fig 3), suggesting that conception-related exposures may exert their influence most prominently during the transition from early to mid-gestation.

**Figure 3.**
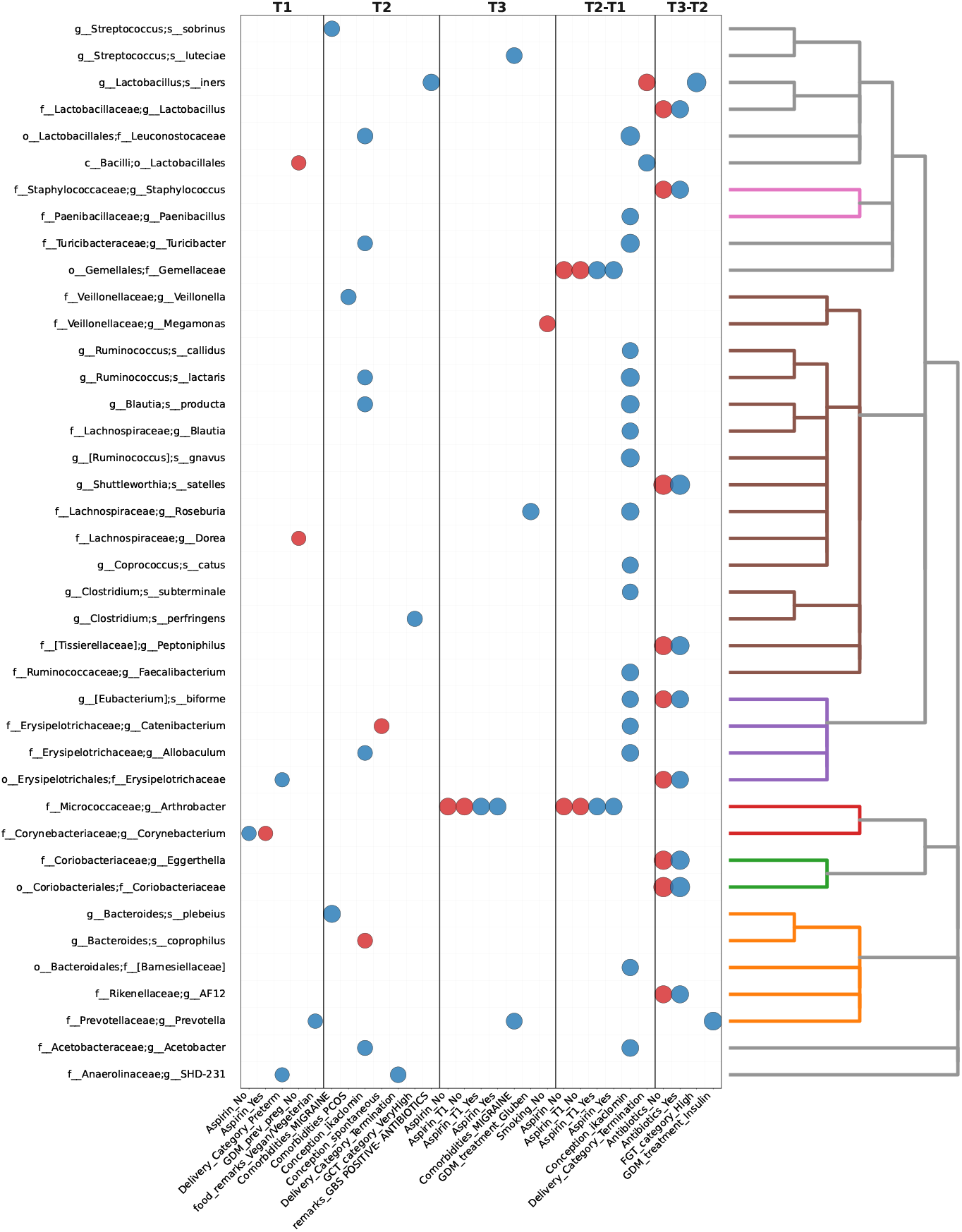
Associations between maternal metadata and microbial taxa. Plot of significant correlations (FDR-corrected, *p* < 0.01) between oral microbiome taxa and maternal metadata across pregnancy trimesters, excluding smoking and gluten-free diet (see Fig. 2). The left y-axis lists microbial taxa, and the right y-axis indicates their phylogeny. The plot is divided into four panels: correlations within T1, within T2, differences between trimesters (T2-T1), and within T3. Differences between T3 and T2 were also analyzed but showed no significant associations after applying FDR correction. Columns represent maternal features (nutrition, demographics, and medical history). Each dot denotes a significant association, with color indicating directionality (blue for positive, red for negative). Dot size is scaled linearly to the absolute correlation coefficient, such that stronger correlations are displayed with larger dots. Spearman’s rank correlation was used for continuous metadata features, and biserial correlation was applied for binary features. See Fig. S19 for the complete heatmap of associations.

To assess whether relationships among maternal features were stable across pregnancy, we computed pairwise correlations among all metadata variables for each trimester (Figs. S26-28). Over-all, a similar network structure of inter-feature dependencies was observed across trimesters, with several features repeatedly forming co-varying clusters. In T1, lifestyle and metabolic variables, such as gluten-free diet, migraine comorbidity, and past smoking were strongly intercorrelated (|*r* |= 0.48-0.54, all *p* < 0.01), while preterm delivery showed negative associations with both Gluten-free diet (*r* = − 0.40, *p* < 0.001) and GCT Very High category (*r* = − 0.31, *p* = 0.046). By T2, this pattern became more pronounced: Gluten-free diet remained correlated with migraine comorbidity (*r* = − 0.52, *p* < 0.001), and a cluster involving smoking (past and current), conception type, and preterm delivery emerged (|*r* |= 0.45-0.77, all *p* < 0.01). These associations suggest increasing interdependence between lifestyle, conception-related, and clinical variables as pregnancy progresses. In T3, inter-feature correlations became sparser, but several consistent relationships persisted. A gluten-free diet remained correlated with past smoking (*r* = 0.52, *p* = 0.008), while migraine comorbidity and antibiotic use formed a minor positive cluster (*r* = 0.31, *p* = 0.007).

To validate the reproducibility of the findings obtained for the Israeli cohort in an independent population, we analyzed the oral microbiome of 154 Russian women sampled during T2 and T3 of pregnancy. PERMANOVA confirmed significant differences in beta diversity between T2 and T3, both for Bray-Curtis dissimilarity (pseudo-F = 2.93, *p* = 0.002) and for the Jaccard index (pseudo-F = 7.10, *p* < 0.001), indicating that temporal compositional shifts were again detectable in this independent population (Figs. S20, S21). At the phylum level, significant decreases in T3 were observed in Euryarchaeota (adjusted *p* = 0.0002), Actinobacteria (adjusted *p* = 0.0487), and Verrucomicrobiota (adjusted *p* = 0.0002), while no phyla exhibited significant increases between trimesters (Fig. S22). Alpha diversity did not change from T2 to T3 (*p* = 0.22). Consistent with the Israeli cohort, GIMIC analysis revealed a pronounced and consistent decrease in Verrucomicrobiota (particularly *A. muciniphila*) and in Methanobacteria across taxonomic levels (see Fig S23). Similarly, miMic analysis identified *A. muciniphila* as significantly reduced from T2 to T3 (adjusted *p* = 0.0005), within genus Veillonella (*V. parvula* and *V. dispar* showed a significant increases in T3 (adjusted *p* = 0.004 for both). As detected in the phylum-level analysis, *Collinsella aerofaciens* (phylum Actinobacteria) showed a consistent decrease in T3 (adjusted *p* = 0.03), while *Prevotella copri* exhibited an even stronger decrease (adjusted *p* < 0.001). *Methanobrevibacter* likewise showed a consistent decrease across all taxonomic levels (adjusted *p* < 0.001) (Fig. S24). Analysis of intra-individual distances further confirmed temporal consistency: cosine distances between samples from the same woman (T2-T3) were significantly smaller than those between different women (*p* < 0.001), supporting subject-specific microbial stability over short time intervals (Fig. S25).

Furthermore, in the Russian cohort, *Dialister* (Veillonellaceae family) in T2 was significantly associated with past smoking history (adjusted *p* = 0.01), aligning with smoking-related associations observed in the Israeli cohort. These results confirm the reproducibility of key microbial shifts and maternal factor associations across both cohorts, highlighting the robustness of the observed oral microbiome dynamics during pregnancy.

## 4 Discussion

We have presented here the first large-scale longitudinal investigation of oral microbiome changes from T1 through T3, addressing a critical gap in understanding early gestational microbial dynamics. Significant changes in the structure of the oral microbial community were observed when comparing T1 to T3. Pairwise comparisons between consecutive trimesters suggest most changes occur in earlier stages of gestation, with stabilization in the T2-T3 period. Although earlier studies have documented oral microbiome alterations, primarily in T2 and T3 [7, **?**, 47], our results show that the most significant changes occur between the first and second trimesters. This pattern aligns with the well-established reduction in microbial richness observed in T3, suggesting a progressive shift toward a distinct microbial community throughout pregnancy. The significant decrease in Verrucomicrobiota, particularly *A. muciniphila*, in the Israeli cohort, and the concurrent increase in Synergistota from T1 to T2 provides important insights into the microbial dynamics of early pregnancy. *A. muciniphila*, a mucin-degrading bacterium extensively studied in the gut microbiome for its roles in maintaining intestinal barrier integrity and metabolic health[5, 39], has also been detected in oral samples, though its prevalence and functional significance in the oral cavity remain less well characterized. The reduction of *A. muciniphila* in the oral cavity during early pregnancy may reflect systemic changes in host-microbe interactions driven by hormonal fluctuations[49, 50].

Synergistota are typically present in low abundance in the oral microbiome, and have been consistently reported in subgingival plaque, periodontal pockets, and endodontic infections, often co-occurring with keystone periodontal pathogens such as *Porphyromonas gingivalis* [26, 6, 43, 49]. Their increased abundance has also been associated with proteolytic metabolism, persistence in inflamed niches, and contribution to chronic biofilm dysbiosis[12]. Thus, the relative increase of Synergistota and the relative decrease in *A. muciniphila* with pregnancy progression may reflect ecological remodeling of the oral microbiome during pregnancy, shifting from a potentially protective profile toward one enriched in taxa linked to tissue inflammation[26].

The increase in Gammaproteobacteria and the decrease in Erysipelotrichia classes further support the concept of a directional shift in the structure of the microbial community between T1 and T2. Gammaproteobacteria are known members of the oral cavity that are often increased under inflammatory or metabolically altered conditions[23]. Erysipelotrichia, in contrast, are more commonly associated with the gut microbiome but have also been detected in saliva[46]. This strengthens the above hypothesis that the oral microbiota in later pregnancy is pro-inflammatory [14, 24].

Despite these alterations, intra-individual profiles remained relatively conserved within individuals compared to between individuals. This suggests that while pregnancy exerts directional pressure on microbial community structure, individual microbial “signatures” persist. The strong association between a gluten-free diet and oral microbiome during pregnancy was among the most intriguing findings in our Israeli cohort. These associations persisted throughout pregnancy. The predominantly positive correlations with taxa within Clostridiales and Bacteroidales, relative to those not adhering to a gluten-free diet. We suggest that eliminating gluten-rich staples may remove competing substrates, enabling expansion of taxa that exploit broader carbohydrate niches. Smoking history was significantly associated with oral community composition across trimesters, with the clearest divergence between those who did and did not previously smoke observed in all trimesters. This interpretation aligns with prior cohort and review evidence showing enrichment of periodontal-associated genera (e.g., *Prevotella, Porphyromonas*) and depletion of commensal Proteobacteria such as *Neisseria*, likely reflecting sustained effects on the oral mucosal environment, including altered oxygen tension, inflammation, and immune modulation [16, 20, 40, 2, 30].

We also identified an association between Ikaclomin conception method and T2 oral microbiome, particularly involving Actinomycetales. Interestingly, this signal appeared to attenuate by T3, suggesting assisted reproductive technologies’ (ART) effects may be strongest in early pregnancy. Possible mechanisms include hormonal stimulation during ART[48] and links between oral dysbiosis and infertility-related conditions like polycystic ovary syndrome or endometriosis[25]. Although the conception method was associated with microbial variation, we did not observe evidence of persistent or long-term effects of ART on maternal microbial health.

These findings were validated in an independent cohort of 154 pregnant women from Russia, underscoring their generalizability across populations and geographic regions. In this cohort, we observed a decline in *A. muciniphila* from T2 to T3, reinforcing its role as a key taxon. In contrast, a number of taxa that had differential abundance throughout pregnancy among Russian women were not duplicated in the Israeli cohort. This may arise from dietary and lifestyle differences or the fact that the majority of the Russian cohort had GDM. Notably, a history of past smoking was again associated with a reduction in *Dialister* abundance at T2, echoing the smoking-related associations found in the Israeli dataset. These results confirm that oral microbiome community dynamics during pregnancy are partially shared across distinct populations, supporting their relevance to maternal-fetal health.

### 4.1 Limitations and Considerations

The observational design precludes causal inference, and oral health status (e.g., gingivitis, caries, salivary pH) was not systematically assessed. Amplicon sequencing was used, restricting taxonomic resolution and functional inference compared with metagenomics or metabolomics. Dietary data were self-reported, raising the possibility of recall bias, and sampling at fixed gestational weeks may not fully capture continuous microbial dynamics. Furthermore, differences in cohort composition (trimesters, GDM prevalence) may affect cross-cohort comparisons.

## 5 Conclusion

Longitudinal profiling across T1-T3 demonstrates pronounced remodeling of the maternal oral microbiota, most evident from T1 to T2, shaped by maternal factors such as smoking history, diet, and mode of conception. Like with the gut microbiota, these shifts in oral microbiota are consistent with a transition toward a pro-inflammatory ecological state. Collectively, we conclude that early-mid pregnancy is a particularly dynamic window in which the oral microbiota may be a relevant therapeutic target. By mapping trimester-specific trajectories tied to modifiable exposures, our work lays the groundwork for microbiota-informed screening and timed intervention in routine obstetrics.

## 6 Declarations

### 6.1 Ethics approval and consent to participate

Informed consent was obtained from all Israeli participants, in accordance with Helsinki ethics permit #0135-15-COM from the Clalit HMO Institutional Review Board and Helsinki ethics permit #0263-15-RMC from the Rabin Medical Center Institutional Review Board. Russian cohort was approved by the local ethics committee of the Almazov National Medical Research Centre, Russia (protocol no. 119)

### 6.2 Availability of data and material

All the datasets are available as taxa-tables with their respective tag in the GitHub repository https://github.com/louzounlab/Early-Pregnancy-Oral-Microbiome. All sequencing data were submitted to EBI (ERP143097).

## Authors’ contributions

S.F. performed the computational analysis and wrote the manuscript. S.T.. contributed data interpretation and writing of the manuscript. P.P., A.S.T., E.A.V., A.D.A., E.A.P., T.M.P., and E.N.G. performed the Russian cohort validation study. Si.F., O.S., A.R., E.T., M.N.O., Y.P., M.H., B.S., and E.H. contributed to data collection and analysis. Y.L. and O.K. supervised this work and wrote the manuscript.

