## Supplementary material for "Early Pregnancy Marks Significant Shifts in the Oral Microbiome": Fig. S*

Table S1: Description of questionnaires and tests included in the study metadata

| **Feature** | **Timepoint** | **Type** | **Description and range** |
|---|---|---|---|
| Medications | T1 | Binary | Chronic medications — Yes / No |
| Aspirin | T1 | Binary | Aspirin during pregnancy — Yes / No |
| Stress | T1, T2, T3 | Categorical | Stress test, 0–4; 0–no stress; 4–high stress |
| Calories | T1, T2, T3 | Continuous | Calories per day (kcal/day) |
| Carb | T1, T2, T3 | Continuous | Carbohydrates per day (g/day or % kcal) |
| Sleeping_hours | T1 | Continuous | Hours per night |
| BMI | Pre-pregnancy | Continuous | $kg/m^2$ |
| BMI_category | Pre-pregnancy | Categorical | Under / Normal / Over / Obese |
| Age | Pre-pregnancy | Continuous | Years (18–43) |
| Age_category | Pre-pregnancy | Categorical | $<25$, 25–34, $\geq 35$ |
| Parity | Pre-pregnancy | Categorical | Number of deliveries |
| Conception | T1 | Categorical | Mode of conception: Spontaneous / Hormonal / IVF / IUI / Ikaclomin |
| Smoking | Pre-pregnancy | Categorical | No / Yes / Past |
| GDM_prev_preg | Pre-pregnancy | Binary | Yes / No |
| Education | Pre-pregnancy | Continuous | Years |
| GCT | T2 | Continuous | Glucose challenge test |
| GCT_category | T2 | Categorical | Glucose challenge test; normal (<140)/ high (140-200)/ very high (>200) |
| FGT | T1 | Continuous | Fasting glucose test |
| FGT_category | T1 | Categorical | Fasting glucose tes; normal (<95), high( 95-126), very high(>126) |
| OGTT_0 | T2/T3 | Binary | Oral glucose tolerance level on time 0 |
| OGTT_60 | T2/T3 | Binary | Oral glucose tolerance test levelafter 60 min |
| OGTT_120 | T2/T3 | Binary | Oral glucose tolerance test level after 120 min |
| OGTT_180 | T2/T3 | Binary | Oral glucose tolerance test level after 180 min |
| OGTT_category | T2/T3 | Categorical | Number of extreme values (0-4); pathologic level- 1-4; normal level- 0 |
| Delivery_week | Post-pregnancy | Continuous | Delivery week |
| Delivery_category | Post-pregnancy | Categorical | < 37- preterm; 37-40- Normal; > 40 - Late; Termination |
| New Born Weight | Post-pregnancy | Continuous | New born weight (kg) |
| GOT | T1 | Continuous | AST; an enzyme found in the liver, elevated levels may indicate liver damage |
| GPT | T1 | Continuous | ALT; an enzyme found in the liver, elevated levels may indicate liver damage |
| PAPP-A | T1 | Continuous | Pregnancy Associated Plasma Protein A; a Placental protein measured in early pregnancy. |

*Continued on next page*

| **Feature** | **Timepoint** | **Type** | **Description** |
|---|---|---|---|
| free_hCG | T1 | Continuous | Human chorionic Gonadotropin; a hormone produced by placenta. |
| AFP | T1 | Continuous | Alpha Fetoprotein, protein produced by the fetal liver, low levels may suggest chromosomal abnormalities. |
| NT | T1 | Continuous | Nuchal Translucency; an ultrasound measurement of fluid at the back of the fetal neck. |
| Extreme0 | T2/T3 | Binary | 1: if > 95 else 0 |
| Extreme60 | T2/T3 | Binary | 1: if > 180 else 0 |
| Extreme120 | T2/T3 | Binary | 1: if > 155 else 0) |
| Extreme80 | T2/T3 | Binary | 1: if > 140 else 0 |
| Antibiotics | T1/T2/T3 | Binary | antibiotics therapy during pregnancy (Yes; No) |
| GDM_treatment | T2/T3 | Category | diet; gluben (oral lower glucose medicine); Insulin (I.M lower glucose medicine) |
| GDM_prev_preg | Pre-pregnancy | Binary | GDM previous pregnancies |
| Medicine | Pre and during pregnancy | Category | Free text |
| Comorbidities | Pre and during pregnancy | Category | Free text |
| food_remarks | Pre-pregnancy | Categorical | Normal / Vegan or Vegetarian / High Carb / Gluten Free |
| Physical_activity | Pre-pregnancy | Continuous | 0–7; 0–low; 7–high |

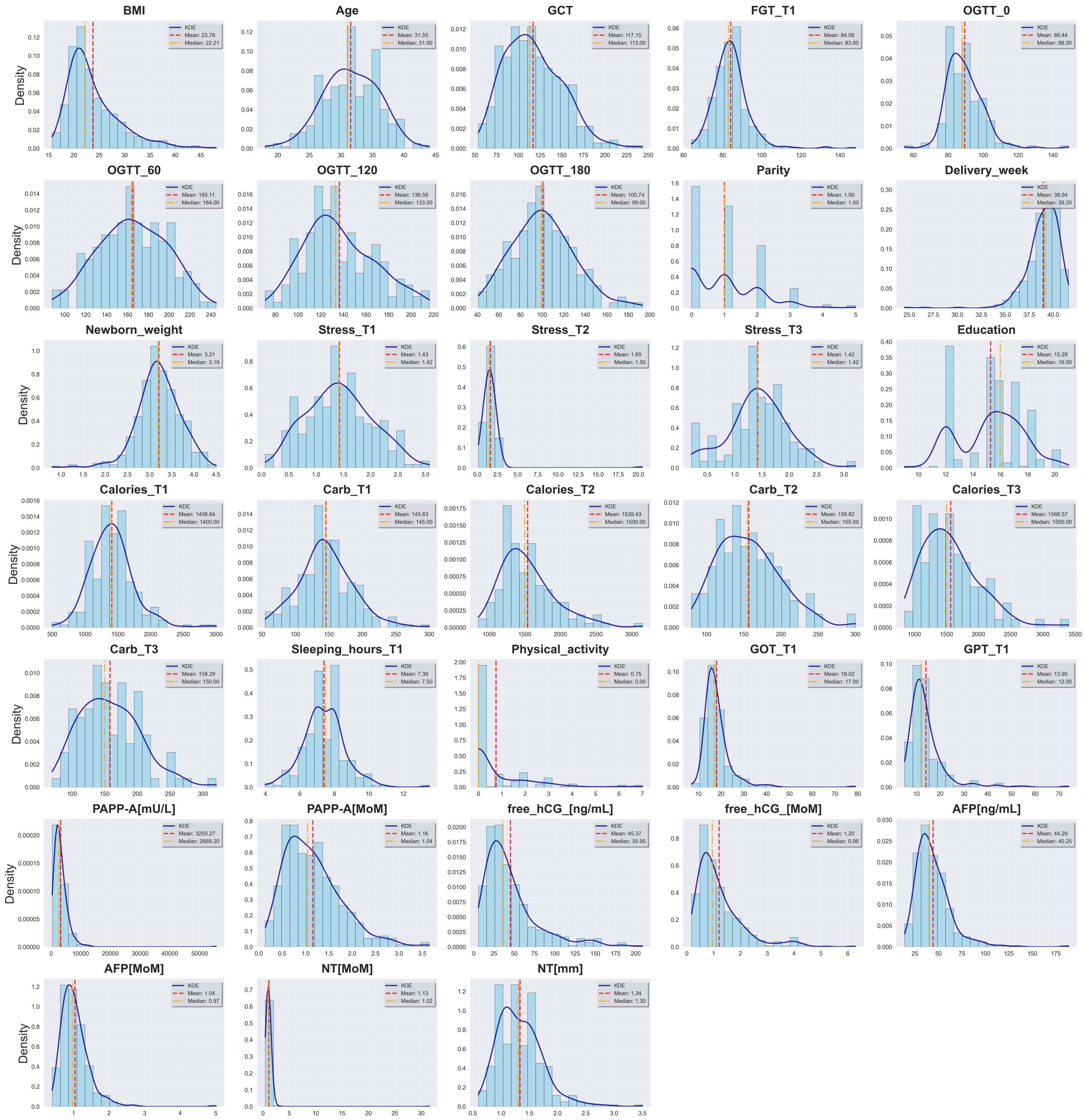


Figure S1: **Feature Distribution Overview.** Comprehensive visualization showing distributions of all Israeli metadata features. (a) Continuous features: Histograms with 20 bins display the data distribution overlaid with kernel density estimation (KDE) curves in dark blue to smooth the distribution pattern. Vertical reference lines indicate the mean (red dashed line) and median (orange dash-dot line) values for each continuous feature.

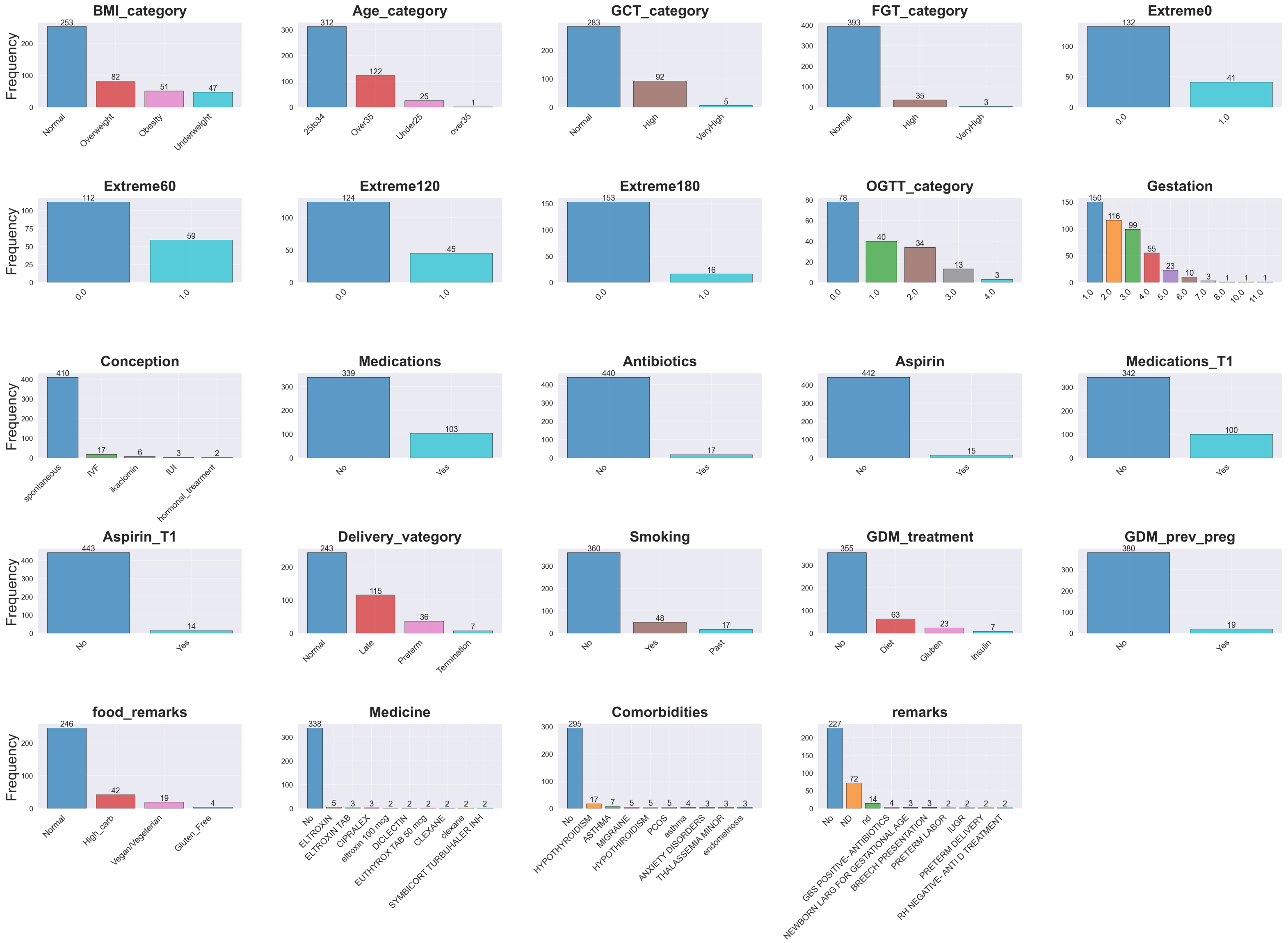


Figure S1: (continued) (b) Categorical and binary features: Bar charts show frequency counts with the actual count labeled on top of each bar for each category within each feature.

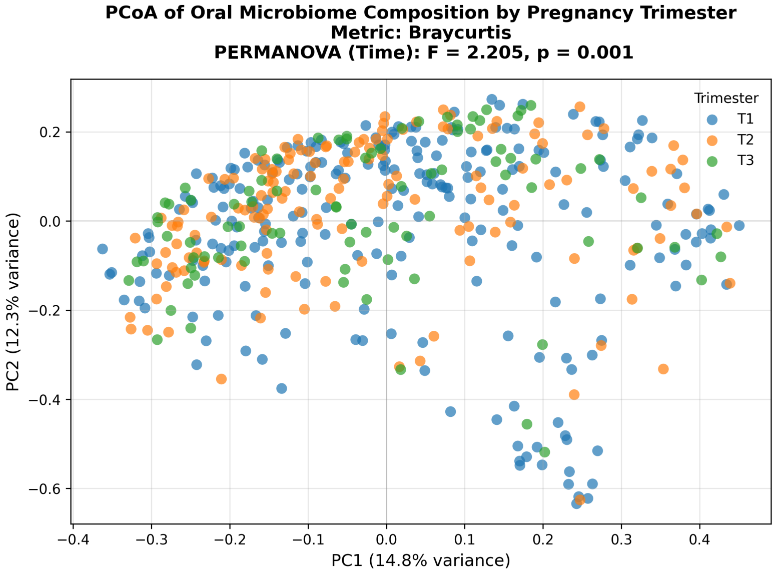


Figure S2: **Principal Coordinates Analysis (PCoA) of Bray-Curtis dissimilarities across pregnancy trimesters.** Each point represents one oral microbiome sample, colored by trimester (T1-T3). The plot is based on relative abundance data. Although clusters overlap in ordination space, PERMANOVA indicated a significant effect of pregnancy stage (pseudo-F = 2.20, $p = 0.0012$), reflecting temporal restructuring of the oral microbial community during gestation.

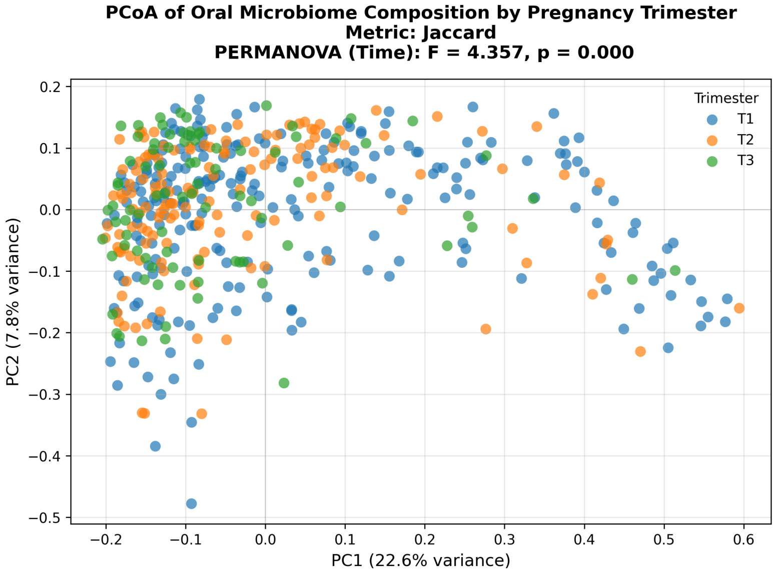


Figure S3: **Principal Coordinates Analysis (PCoA) of Jaccard dissimilarities across pregnancy trimesters.** PCoA based on presence-absence profiles, colored by trimester (T1-T3). PERMANOVA confirmed a significant temporal effect on microbial community composition (pseudo-F $= 2.31$, $p < 0.01$), consistent with dynamic changes in oral microbiome structure during pregnancy.

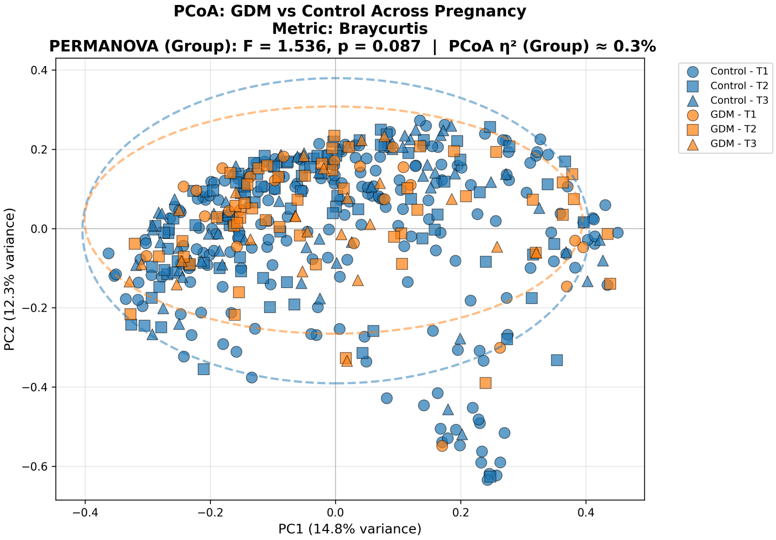


Figure S4: **Principal Coordinates Analysis (PCoA) of Bray–Curtis dissimilarities comparing GDM and control samples.** Points represent samples colored by group (GDM = orange, Control = blue) and shaped by trimester (T1–T3). No distinct separation was observed between GDM and control samples (PERMANOVA pseudo-F $= 1.54$, $p = 0.092$; $\eta^2 = 0.3$ %), indicating highly similar microbial community structures across groups.

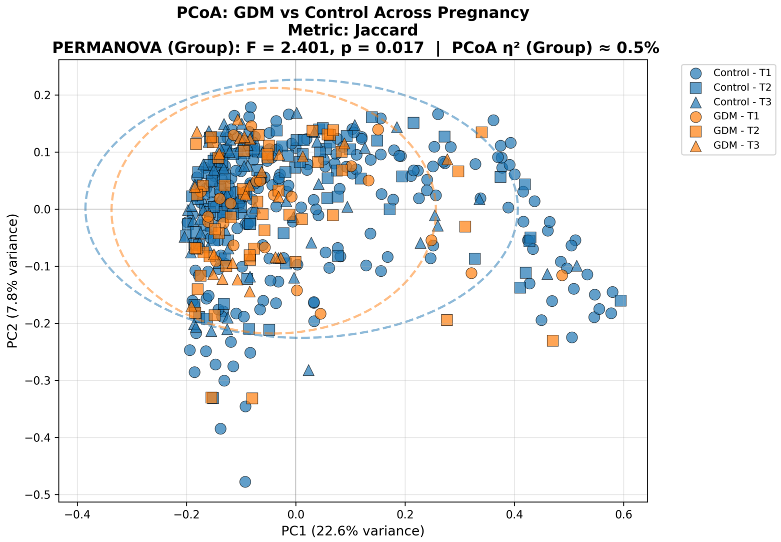


Figure S5: **Principal Coordinates Analysis (PCoA) of Jaccard dissimilarities comparing GDM and control samples.** PCoA based on presence–absence data, colored by GDM status and shaped by trimester. A small but statistically significant group effect was detected (PERMANOVA pseudo-F = 2.40, $p = 0.014$, FDR-adjusted $p = 0.030$), although the variance explained was minimal ($\eta^2 = 0.5$ %), indicating that GDM status contributes little to overall community composition. No significant GDM–control differences were observed within individual trimesters (all FDR-adjusted p ¿ 0.05; see Table **??**).

Table S2: Summary statistics (mean and median) of microbial phyla across trimesters

| **Phyla** | **Mean; Median T1** | **Mean; Median T2** | **Mean; Median T3** |
|---|---|---|---|
| Euryarchaeota | 3.20; 0 | 1.22 ; 0 | 1.17 ; 0 |
| Actinobacteria | 15.31 ; 13.11 | 13.40 ; 11.68 | 15.95 ; 14.61 |
| Bacteroidetes | 10.61 ; 9.00 | 10.05 ; 9.24 | 9.32 ; 7.28 |
| Chloroflexi | 0.15 ; 0 | 0.08 ; 0 | 0.98 ; 0 |
| Cyanobacteria | 0.57 ; 0 | 0.12 ; 0 | 0.02 ; 0 |
| Deferribacteres | 1.02 ; 0 | 0.34 ; 0 | 0.14 ; 0 |
| Firmicutes | 12.74 ; 11.33 | 13.85 ; 12.99 | 14.28 ; 13.66 |
| Fusobacteria | 11.19 ; 10.46 | 13.90 ; 11.07 | 12.90 ; 11.15 |
| Proteobacteria | 18.29 ; 14.62 | 20.43 ; 17.27 | 24.01 ; 21.31 |
| SR1 | 5.18 ; 1.26 | 4.68 ; 1.14 | 4.58 ; 1.44 |
| Spirochaetes | 3.72 ; 2.53 | 3.85 ; 2.82 | 4.37 ; 2.78 |
| Synergistetes | 6.12 ; 2.93 | 11.11 ; 5.64 | 7.32 ; 4.26 |
| TM7 | 0.69 ; 0 | 0.96 ; 0 | 1.27 ; 0 |
| Tenericutes | 0.72 ; 0.20 | 0.54 ; 0.18 | 0.49 ; 0.11 |
| Verrucomicrobia | 10.42 ; 1.28 | 5.40 ; 0.51 | 3.13 ; 0.50 |

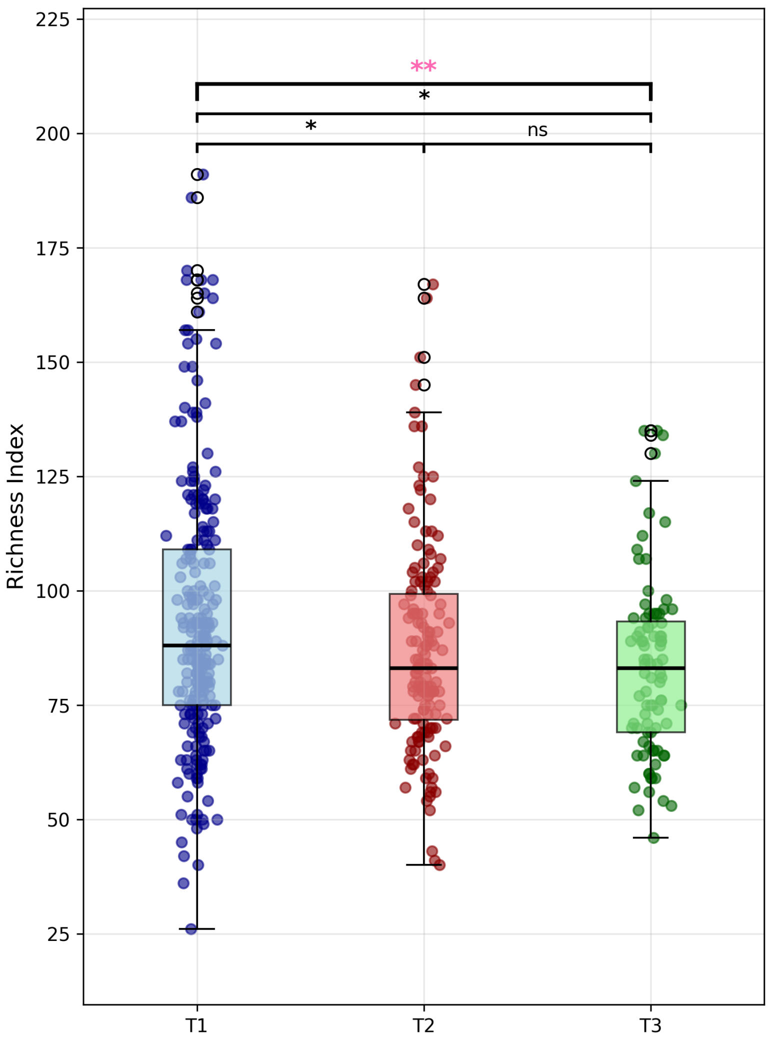


Figure S6: **Comparison of alpha diversity between pregnancy trimesters.** Boxplots show the distribution of species richness based on relative abundance data. Statistical significance was assessed using the Kruskal–Wallis test ($p = 0.0064$, shown in pink), followed by pairwise Mann–Whitney U tests. A significant decrease was observed from T1 to T2 (adjusted $p = 0.046$) and from T1 to T3 (adjusted $p = 0.012$), whereas the decrease from T2 to T3 was not significant (adjusted $p = 0.264$).

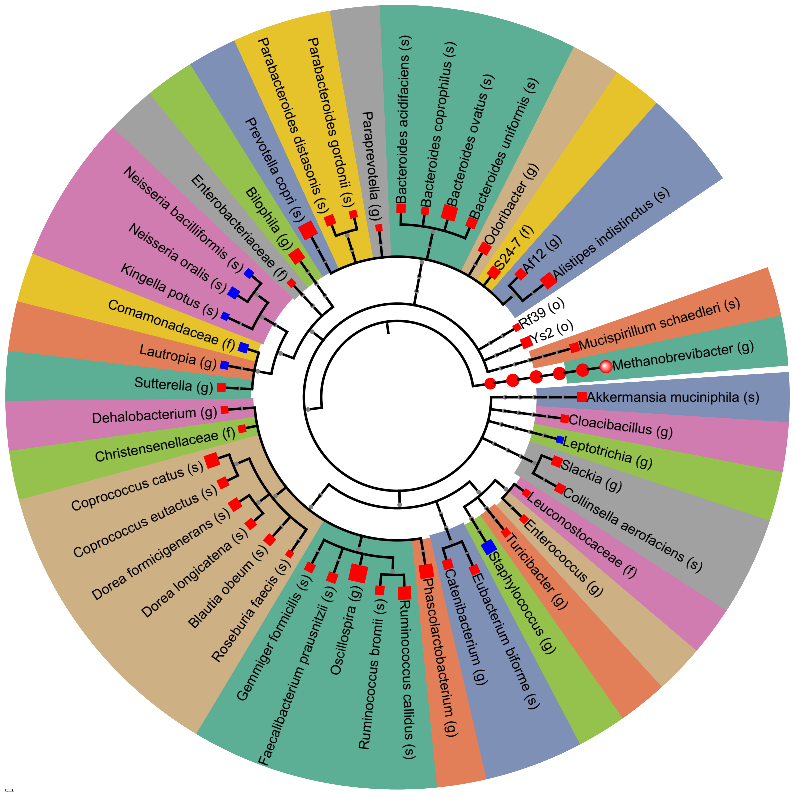


Figure S7: miMic test, identifying microbes with significant differences between T1 and T3, where each significant taxon is mapped onto the taxonomic tree. Dot color indicates direction (blue = increased in T3, red = decreased in T3), and dot size reflects the absolute Mann–Whitney statistic coefficient. The background shading groups taxa that belong to the same family.

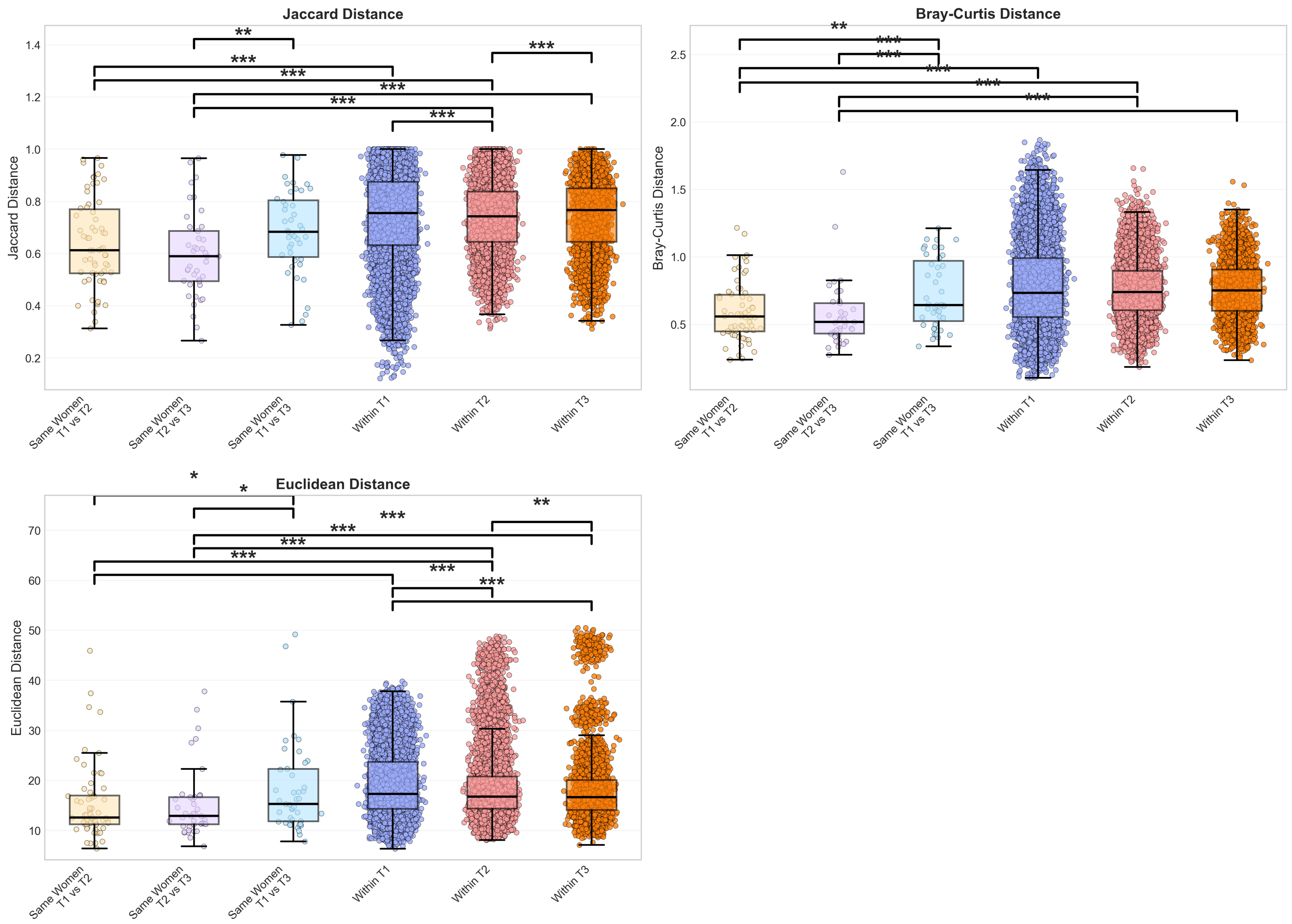


Figure S8: **Distance boxplots across multiple metrics.** Boxplots of distance metrics by comparison type: same women across trimesters, different women within trimesters, and different women between trimesters. Higher values indicate greater dissimilarity. Asterisks denote significance ($*p < 0.05$, $**p < 0.01$, $***p < 0.001$). Same-women distances are consistently smaller. Comparisons were made using the Mann-Whitney U test.

### 0.1 Circular taxonomic tree mapping microbes significantly associated with maternal characteristics in oral microbiome

Circular taxonomic trees showing microbial taxa significantly associated with specific maternal features in a specific trimester. Each tree follows the same format as the miMic tree in Fig. 1E. Dot colors indicate directionality of association (blue for positive association, red for negative association), dot size reflects the magnitude of the association, and background shading groups taxa by family. The trees are filtered to show only significant genus and species taxa.

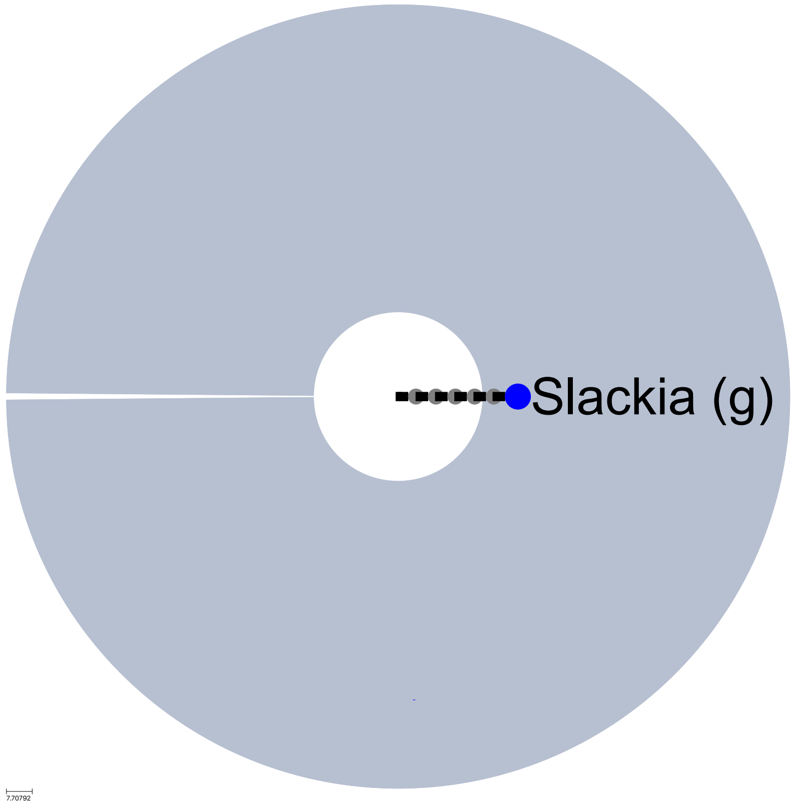


Figure S9: **Gluten-free diet in first trimester:** Blue indicates increased abundance in women following a gluten-free diet; red indicates decreased abundance.

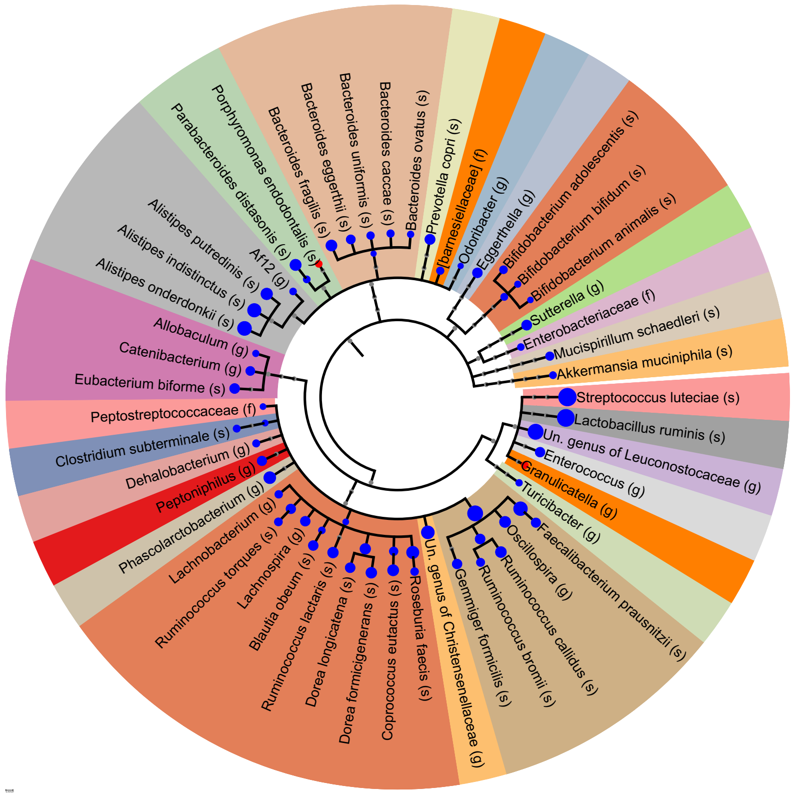


Figure S10: **Gluten-free diet in second trimester:** Blue indicates increased abundance in women following a gluten-free diet; red indicates decreased abundance.

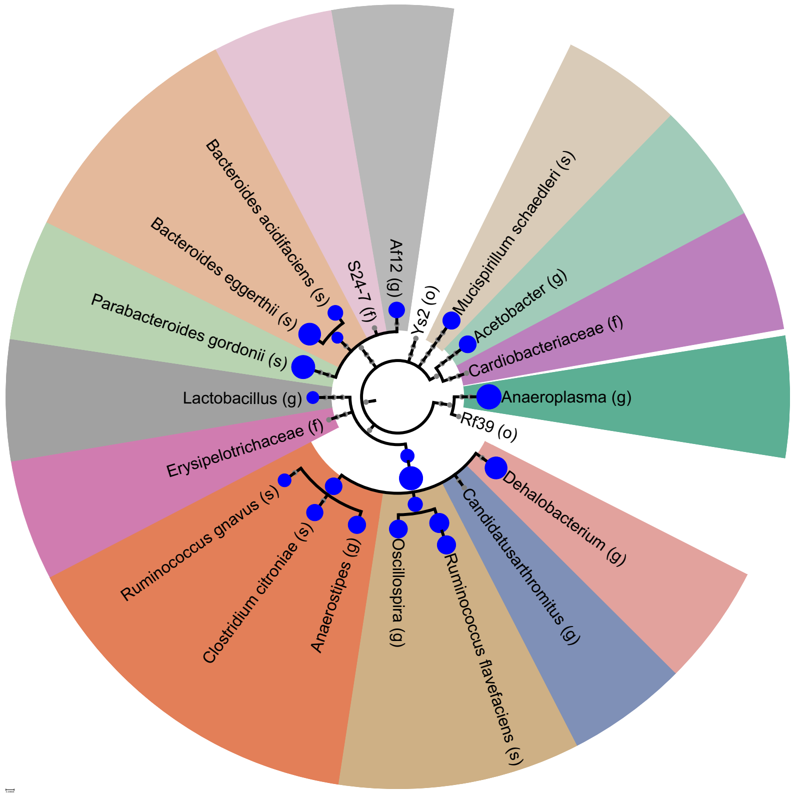


Figure S11: **Gluten-free diet in third trimester:** Blue indicates increased abundance in women following a gluten-free diet; red indicates decreased abundance.

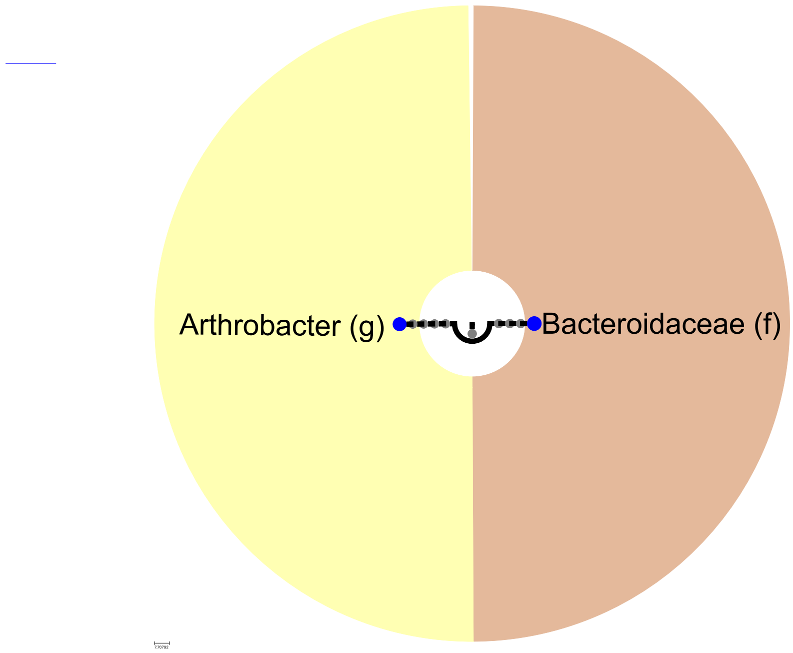


Figure S12: **Smoking history in first trimester:** Blue indicates increased abundance in women who smoked in the past; red indicates decreased abundance.

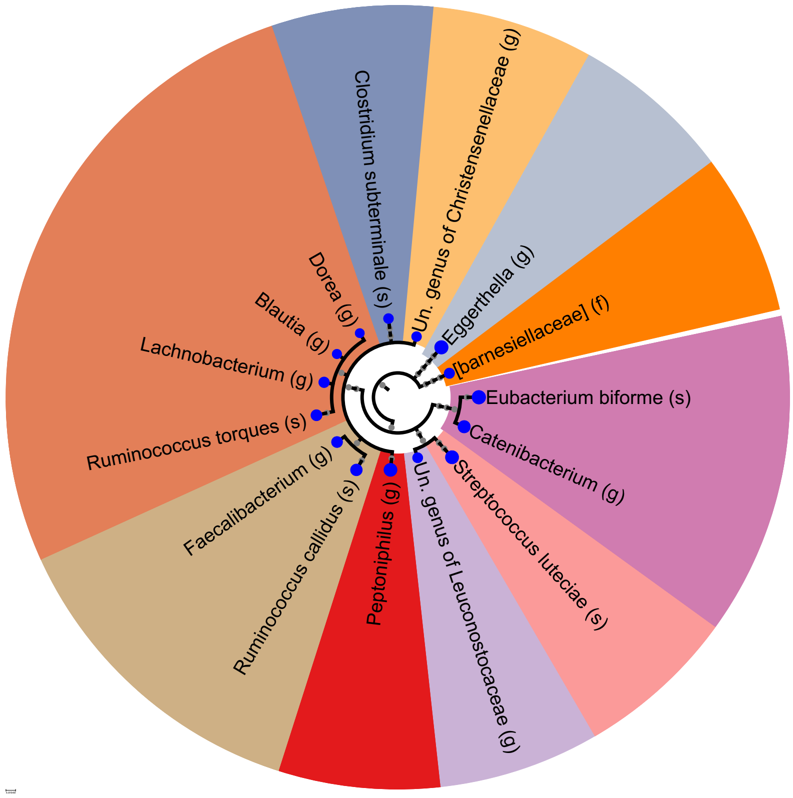


Figure S13: **Smoking history in second trimester:** Blue indicates increased abundance in women who smoked in the past; red indicates decreased abundance.

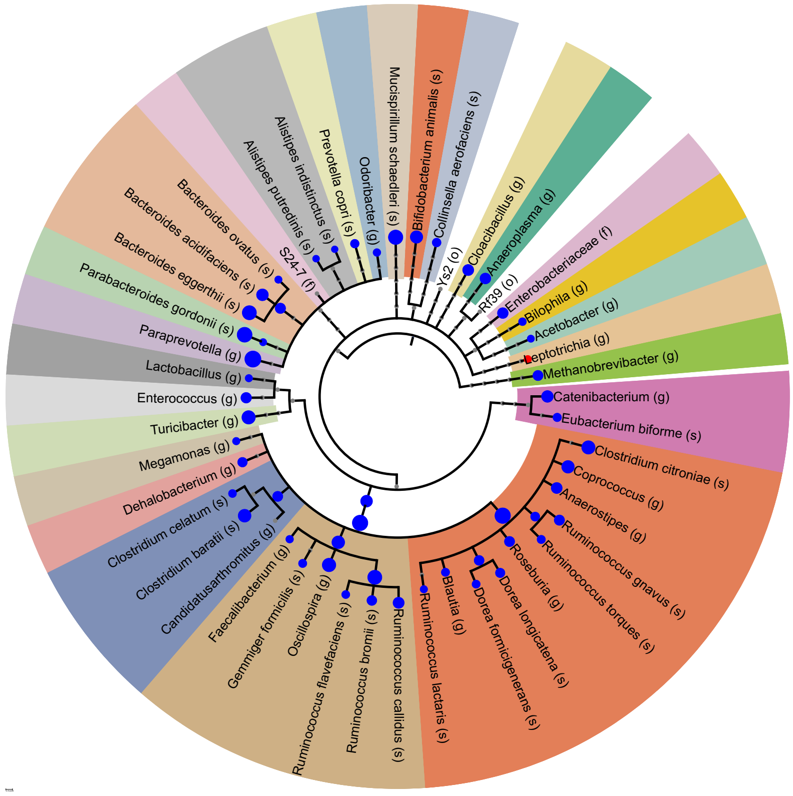


Figure S14: **Smoking history in third trimester:** Blue indicates increased abundance in women who smoked in the past; red indicates decreased abundance.

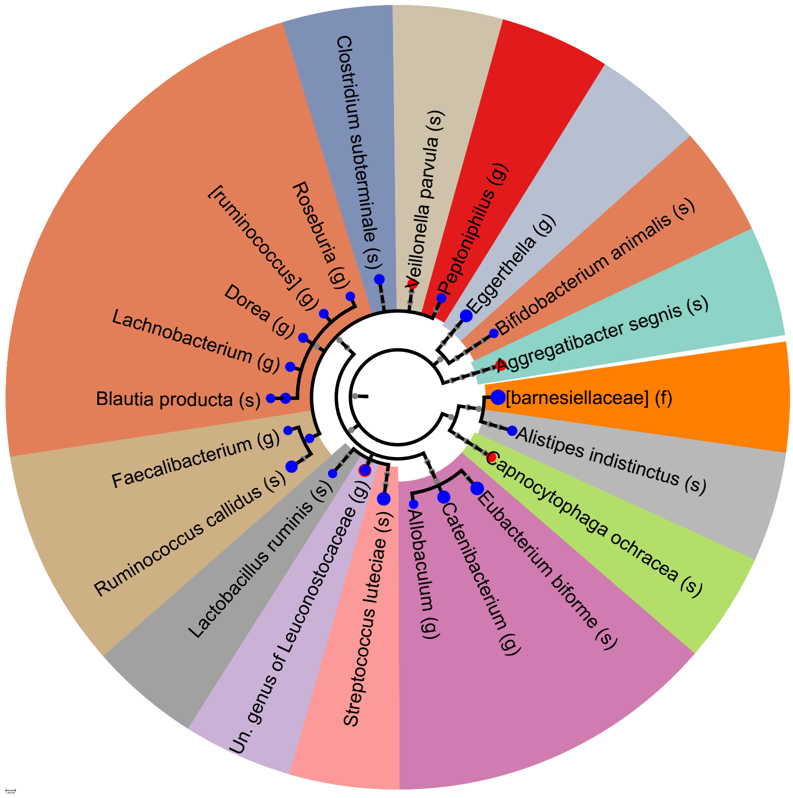


Figure S15: **Conception with ikaclomin in second trimester:** Blue indicates increased abundance in women who conceived with ikaclomin; red indicates decreased abundance.

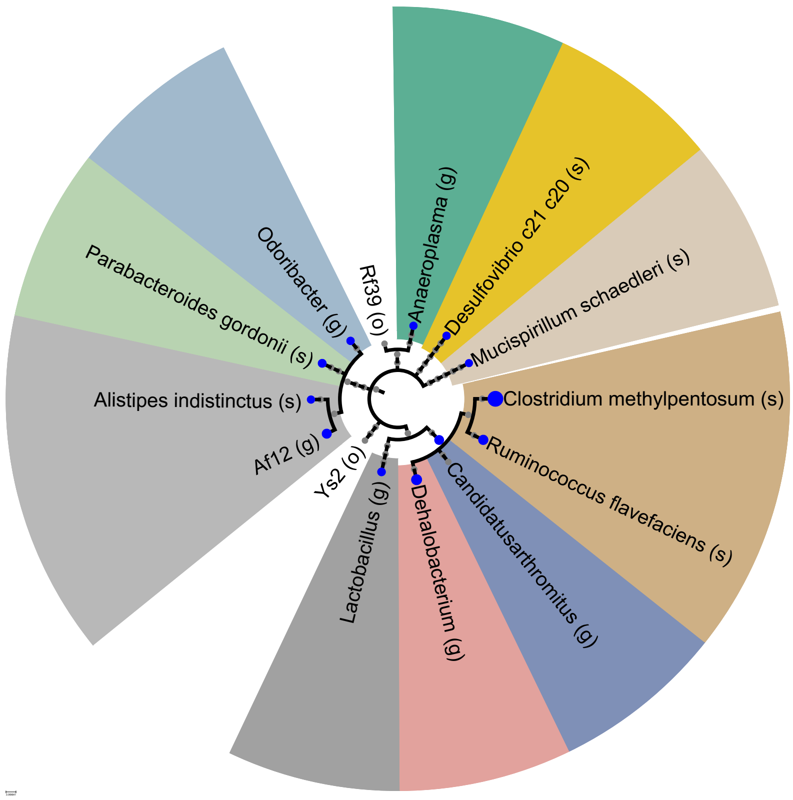


Figure S16: **Conception with hormonal treatment in first trimester:** Blue indicates increased abundance in women who conceived with hormonal treatment; red indicates decreased abundance.

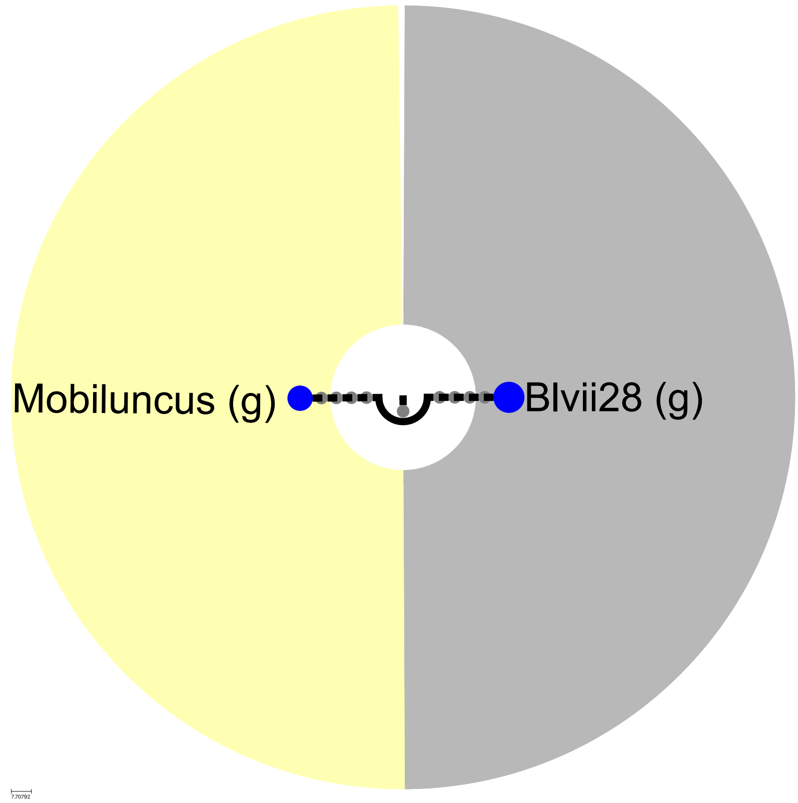


Figure S17: **Conception with hormonal treatment in the second trimester:** Blue indicates increased abundance in women who conceived with hormonal treatment; red indicates decreased abundance.

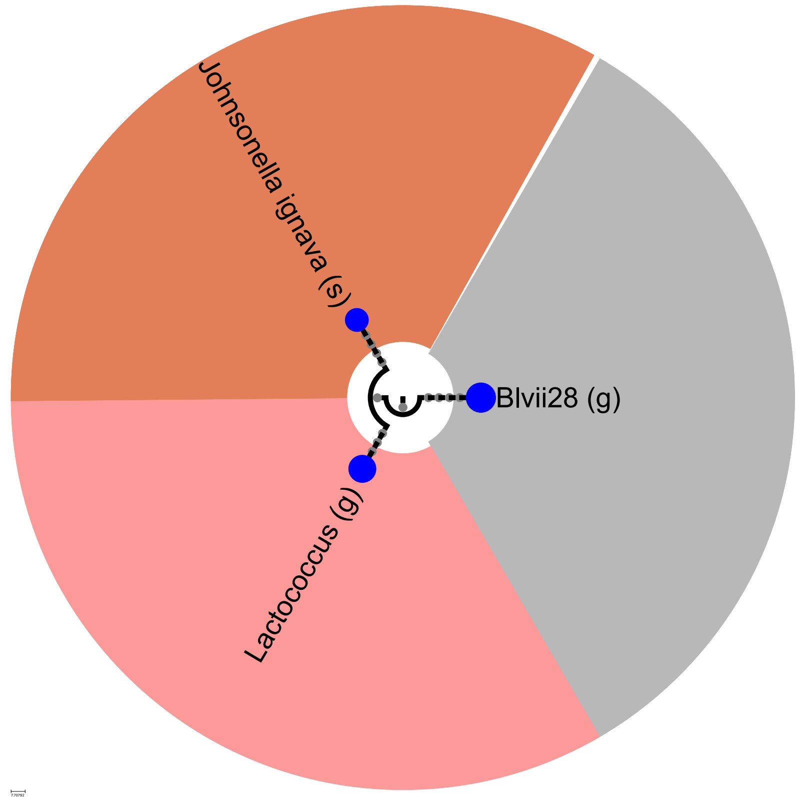


Figure S18: **Conception with hormonal treatment in third trimester:** Blue indicates increased abundance in women who conceived with hormonal treatment; red indicates decreased abundance.

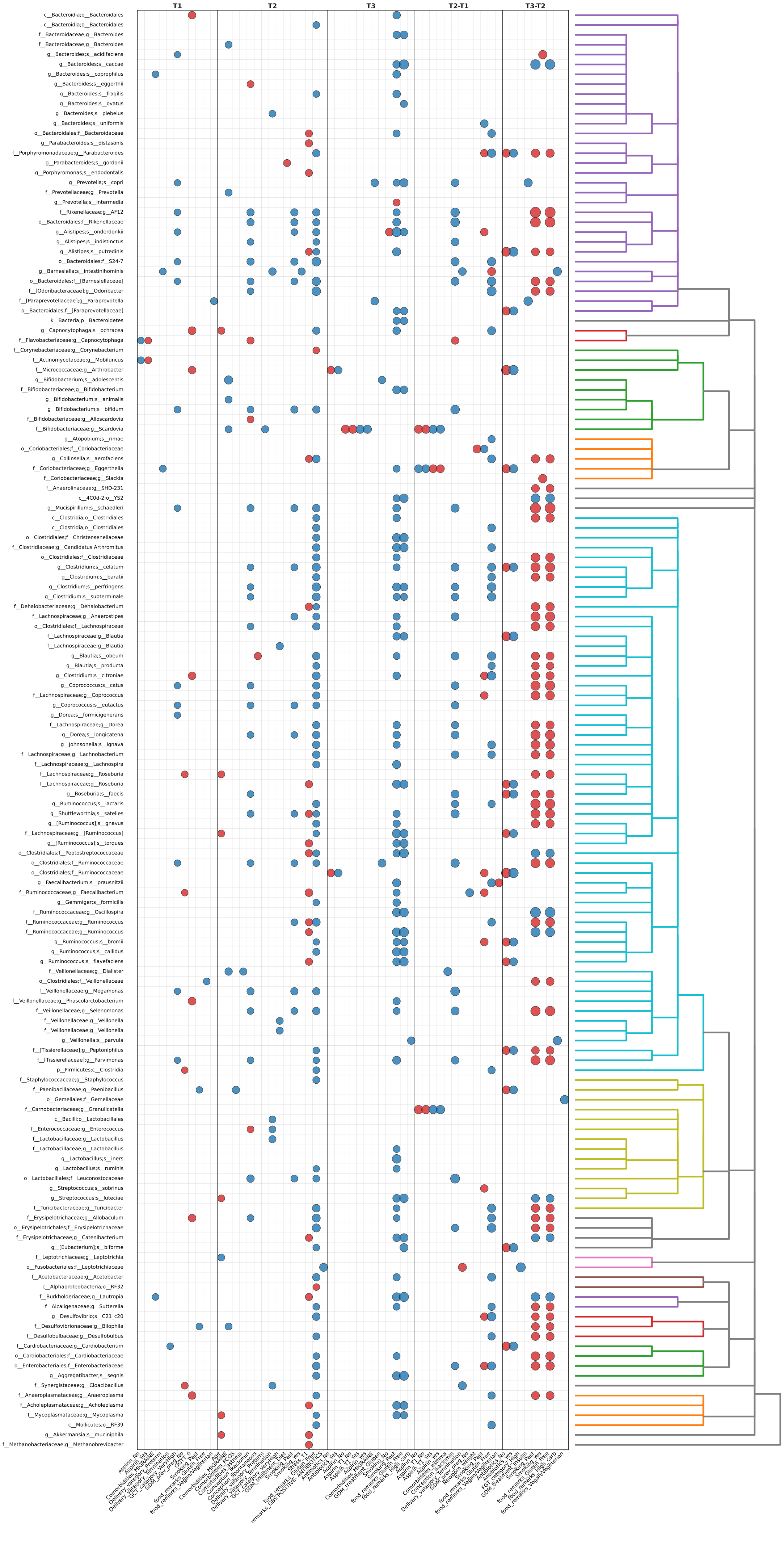
T1
T2
T3
T2-T1
T3-T2
c__Bacteroidia;o__Bacteroidales
c__Bacteroidia;o__Bacteroidales
f__Bacteroidaceae;g__Bacteroides
f__Bacteroidaceae;g__Bacteroides
g__Bacteroides;s__acidifaciens
g__Bacteroides;s__caccae
g__Bacteroides;s__coprophilus
g__Bacteroides;s__eggerthii
g__Bacteroides;s__fragilis
g__Bacteroides;s__ovatus
g__Bacteroides;s__plebeius
g__Bacteroides;s__uniformis
o__Bacteroidales;f__Bacteroidaceae
g__Parabacteroides;s__distasonis
f__Porphyromonadaceae;g__Parabacteroides
g__Parabacteroides;s__gordonii
g__Porphyromonas;s__endodontalis
g__Prevotella;s__copri
f__Prevotellaceae;g__Prevotella
g__Prevotella;s__intermedia
f__Rikenellaceae;g__AF12
o__Bacteroidales;f__Rikenellaceae
g__Alistipes;s__onderdonkii
g__Alistipes;s__indistinctus
g__Alistipes;s__putredinis
o__Bacteroidales;f__S24-7
g__Barnesiella;s__intestinihominis
o__Bacteroidales;f__[Barnesiellaceae]
f__[Odoribacteraceae];g__Odoribacter
f__[Paraprevotellaceae];g__Paraprevotella
o__Bacteroidales;f__[Paraprevotellaceae]
k__Bacteria;p__Bacteroidetes
g__Capnocytophaga;s__ochracea
f__Flavobacteriaceae;g__Capnocytophaga
f__Corynebacteriaceae;g__Corynebacterium
f__Actinomycetaceae;g__Mobiluncus
f__Micrococcaceae;g__Arthrobacter
g__Bifidobacterium;s__adolescentis
f__Bifidobacteriaceae;g__Bifidobacterium
g__Bifidobacterium;s__animalis
g__Bifidobacterium;s__bifidum
f__Bifidobacteriaceae;g__Alloscardovia
f__Bifidobacteriaceae;g__Scardovia
g__Atopobium;s__rimae
o__Coriobacteriales;f__Coriobacteriaceae
g__Collinsella;s__aerofaciens
f__Coriobacteriaceae;g__Eggerthella
f__Coriobacteriaceae;g__Slackia
f__Anaerolinaceae;g__SHD-231
c__4C0d-2;o__YS2
g__Mucispirillum;s__schaedleri
c__Clostridia;o__Clostridiales
c__Clostridia;o__Clostridiales
o__Clostridiales;f__Christensenellaceae
f__Clostridiaceae;g__Candidatus Arthromitus
o__Clostridiales;f__Clostridiaceae
g__Clostridium;s__celatum
g__Clostridium;s__baratii
g__Clostridium;s__perfringens
g__Clostridium;s__subterminale
f__Dehalobacteriaceae;g__Dehalobacterium
f__Lachnospiraceae;g__Anaerostipes
o__Clostridiales;f__Lachnospiraceae
f__Lachnospiraceae;g__Blautia
f__Lachnospiraceae;g__Blautia
g__Blautia;s__obeum
g__Blautia;s__producta
g__Clostridium;s__citroniae
g__Coprococcus;s__catus
f__Lachnospiraceae;g__Coprococcus
g__Coprococcus;s__eutactus
g__Dorea;s__formicigenerans
f__Lachnospiraceae;g__Dorea
g__Dorea;s__longicatena
g__Johnsonella;s__ignava
f__Lachnospiraceae;g__Lachnobacterium
f__Lachnospiraceae;g__Lachnospira
f__Lachnospiraceae;g__Roseburia
f__Lachnospiraceae;g__Roseburia
g__Roseburia;s__faecis
g__Ruminococcus;s__lactaris
g__Shuttleworthia;s__satelles
g__[Ruminococcus];s__gnavus
f__Lachnospiraceae;g__[Ruminococcus]
g__[Ruminococcus];s__torques
o__Clostridiales;f__Peptostreptococcaceae
o__Clostridiales;f__Ruminococcaceae
o__Clostridiales;f__Ruminococcaceae
g__Faecalibacterium;s__prausnitzii
f__Ruminococcaceae;g__Faecalibacterium
g__Gemmiger;s__formicilis
f__Ruminococcaceae;g__Oscillospira
f__Ruminococcaceae;g__Ruminococcus
f__Ruminococcaceae;g__Ruminococcus
g__Ruminococcus;s__bromii
g__Ruminococcus;s__callidus
g__Ruminococcus;s__flavefaciens
f__Veillonellaceae;g__Dialister
o__Clostridiales;f__Veillonellaceae
f__Veillonellaceae;g__Megamonas
f__Veillonellaceae;g__Phascolarctobacterium
f__Veillonellaceae;g__Selenomonas
f__Veillonellaceae;g__Veillonella
f__Veillonellaceae;g__Veillonella
g__Veillonella;s__parvula
f__[Tissierellaceae];g__Peptoniphilus
f__[Tissierellaceae];g__Parvimonas
p__Firmicutes;c__Clostridia
f__Staphylococcaceae;g__Staphylococcus
f__Paenibacillaceae;g__Paenibacillus
o__Gemellales;f__Gemellaceae
f__Carnobacteriaceae;g__Granulicatella
c__Bacilli;o__Lactobacillales
f__Enterococcaceae;g__Enterococcus
f__Lactobacillaceae;g__Lactobacillus
f__Lactobacillaceae;g__Lactobacillus
g__Lactobacillus;s__iners
g__Lactobacillus;s__ruminis
o__Lactobacillales;f__Leuconostocaceae
g__Streptococcus;s__sobrinus
g__Streptococcus;s__luteciae
f__Turicibacteraceae;g__Turicibacter
f__Erysipelotrichaceae;g__Allobaculum
o__Erysipelotrichales;f__Erysipelotrichaceae
f__Erysipelotrichaceae;g__Catenibacterium
g__[Eubacterium];s__biforme
f__Leptotrichiaceae;g__Leptotrichia
o__Fusobacteriales;f__Leptotrichiaceae
f__Acetobacteraceae;g__Acetobacter
c__Alphaproteobacteria;o__RF32
f__Burkholderiaceae;g__Lautropia
f__Alcaligenaceae;g__Sutterella
g__Desulfovibrio;s__C21_c20
f__Desulfovibrionaceae;g__Bilophila
f__Desulfobulbaceae;g__Desulfobulbus
f__Cardiobacteriaceae;g__Cardiobacterium
o__Cardiobacteriales;f__Cardiobacteriaceae
o__Enterobacteriales;f__Enterobacteriaceae
g__Aggregatibacter;s__segnis
f__Synergistaceae;g__Cloacibacillus
f__Anaeroplasmataceae;g__Anaeroplasma
f__Acholeplasmataceae;g__Acholeplasma
f__Mycoplasmataceae;g__Mycoplasma
c__Mollicutes;o__RF39
g__Akkermansia;s__muciniphila
f__Methanobacteriaceae;g__Methanobrevibacter

Figure S19: **Associations between maternal metadata and microbial taxa.** Heatmap of significant correlations (FDR-corrected, $p < 0.05$) between oral microbiome taxa and maternal metadata across pregnancy trimesters. The left y-axis lists microbial taxa, and the right y-axis indicates their taxonomic linkage. The heatmap is divided into four panels: correlations within T1, within T2, within T3, and differences between trimesters (T2–T1 and T3-T2). Columns represent maternal features (nutrition, demographics, and medical history) with significant associations (FDR-corrected, $p < 0.05$). Each dot denotes a significant association, with color indicating directionality (blue for positive, red for negative) and size reflecting the absolute value of the test statistic. Spearman's rank correlation was used for continuous metadata features, and biserial correlation was applied for binary features.

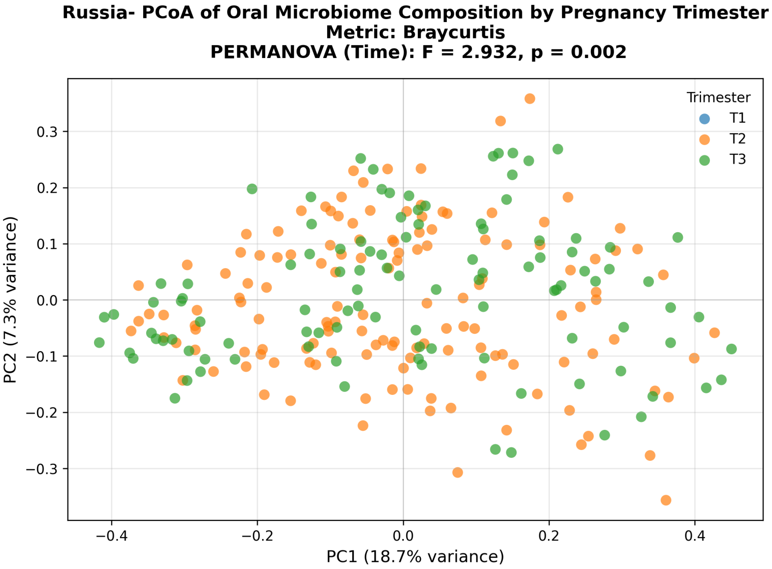


Figure S20: **Principal Coordinates Analysis (PCoA) of Bray–Curtis dissimilarities across pregnancy trimesters- Russian cohort.** Each point represents one oral microbiome sample, colored by trimester (only T2 and T3 available in the Russian cohort). The plot is based on relative abundance data. Although clusters overlap in ordination space, PERMANOVA indicated a significant effect of pregnancy stage (pseudo-F = 2.93, $p = 0.002$), reflecting temporal restructuring of the oral microbial community during gestation.

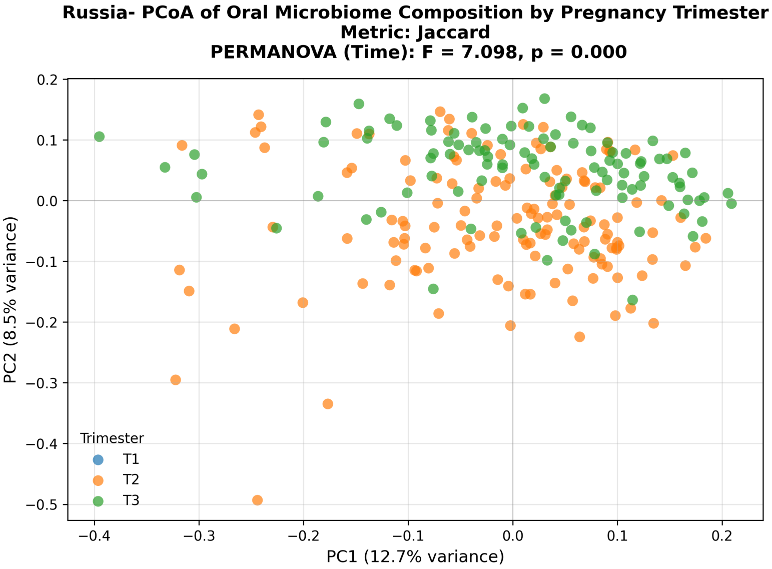


Figure S21: **Principal Coordinates Analysis (PCoA) of Jaccard dissimilarities across pregnancy trimesters- Russian cohort.** PCoA based on presence–absence profiles, colored by trimester (only T2 and T3 available in the Russian cohort). PERMANOVA confirmed a significant temporal effect on microbial community composition (pseudo-F = 7.09, $p < 0.001$), consistent with dynamic changes in oral microbiome structure during pregnancy.

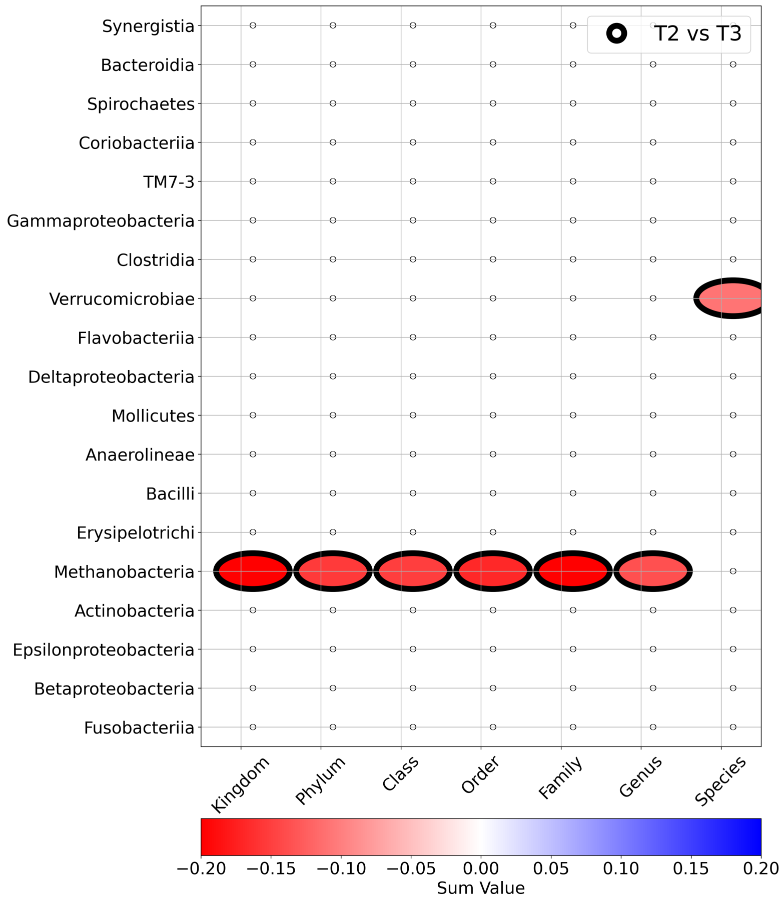


Figure S23: GIMIC results of Russia cohort, depicting a consistent taxonomic shift with Israeli cohort, with edge colors indicating comparisons: black for T2 vs. T3. Bubbles summarize total differences across trimesters at each taxonomic level, with colors indicating enrichment (T2 vs. T3: red for T2, blue for T3), highlighting consistent shifts across taxonomic levels.

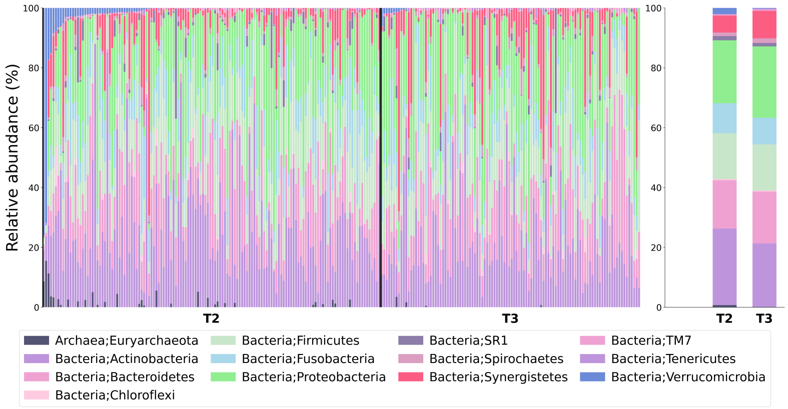


Figure S22: Stacked bar plots of Russia cohort microbial composition at the phylum level, with each color representing a different phylum and each bar corresponding to one sample (summing to 100% relative abundance per sample). The average distribution on the right highlights a decrease in Verrucomicrobiota during pregnancy.

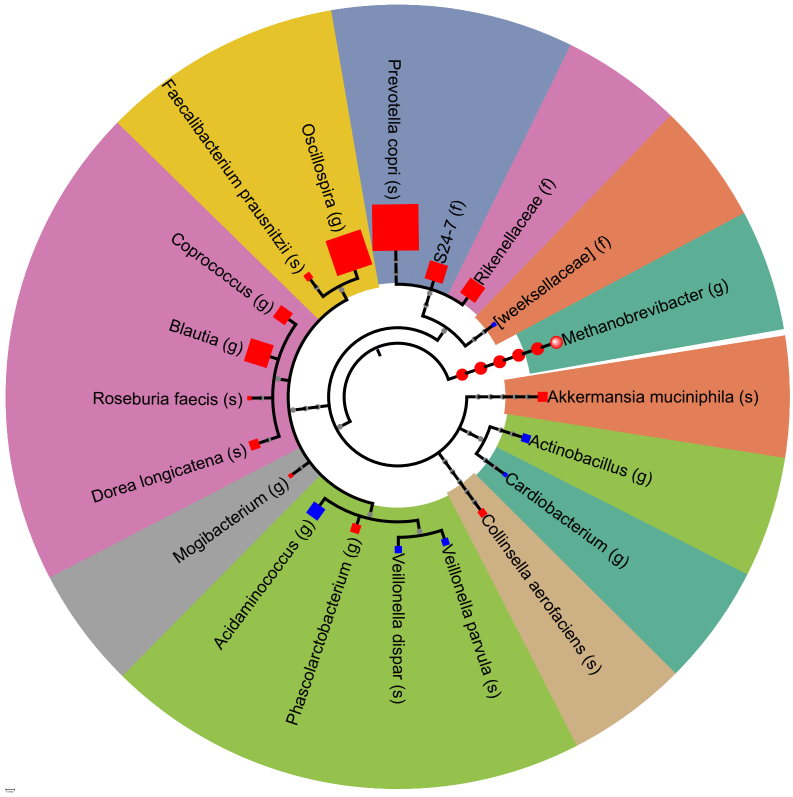


Figure S24: miMic test, identifying microbes with significant differences between T2 and T3 in the Russian cohort, where each significant taxon is mapped onto the taxonomic tree. Dot color indicates direction (blue = increased in T3, red = decreased in T3), and dot size reflects the absolute Mann–Whitney statistic coefficient. The background shading groups taxa that belong to the same family.

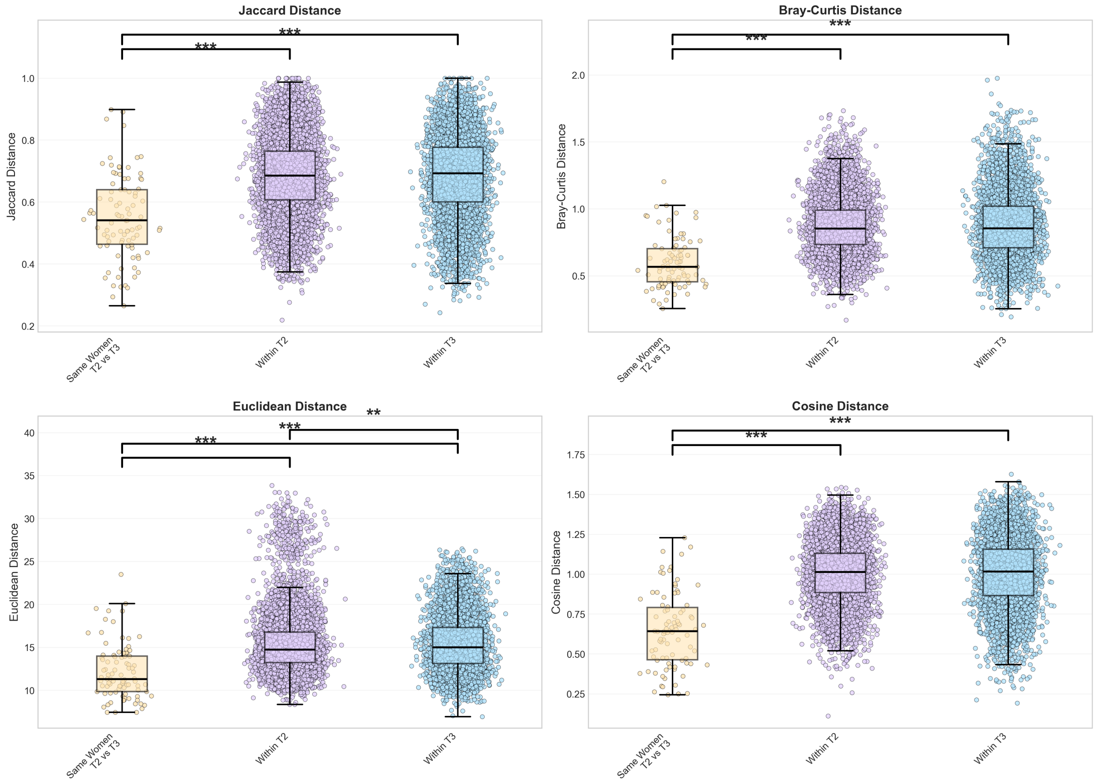


Figure S25: Jaccard, Bray-Curtis, Euclidean, and Cosine distance metrics boxplots comparing microbiome dissimilarity across scenarios, with colors distinguishing comparison types: same women across trimesters, different women within trimesters, and different women between trimesters. Higher values indicate greater dissimilarity, with significance marked by asterisks ($*p < 0.05$, $**p < 0.01$, $***p < 0.001$). Distances between samples from the same woman were consistently smaller than in other scenarios.

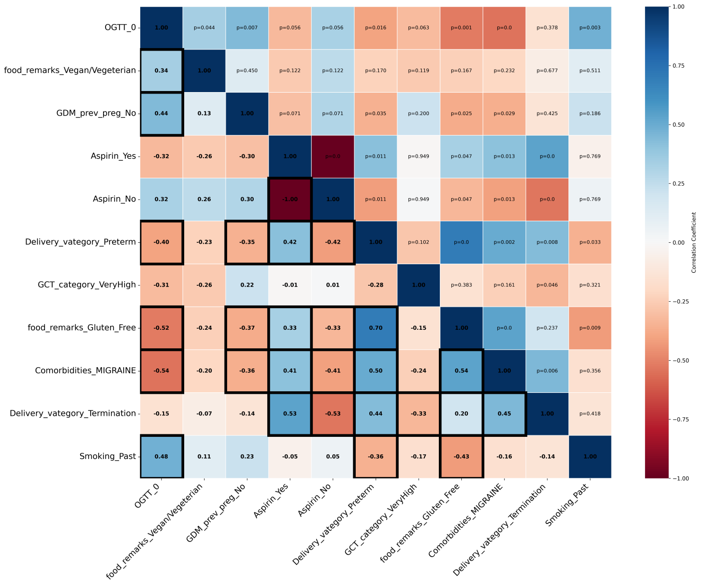


Figure S26: Correlation matrix of Israeli maternal features during the first trimester (T1). Lower triangle: Pearson correlation coefficients; upper triangle: p-values; black edges denote $p < 0.05$.

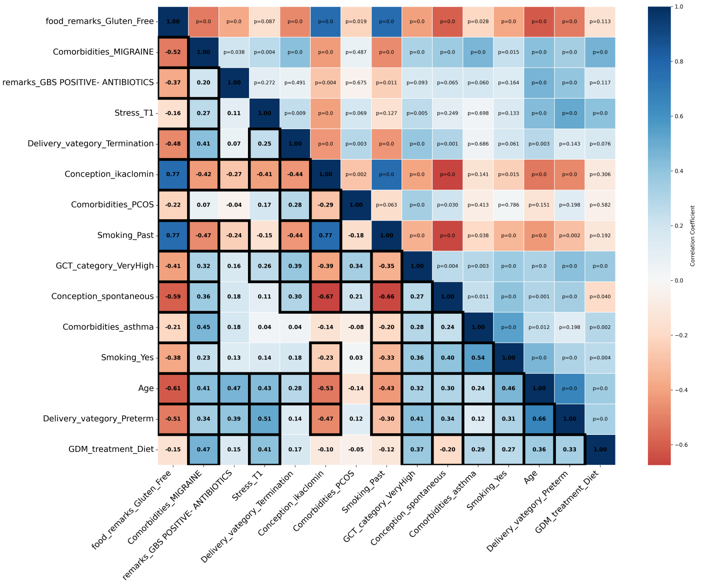


Figure S27: Correlation matrix of Israeli maternal features during the second trimester (T2). A similar structure was observed as in T1, with stronger clustering among lifestyle and conception-related variables.

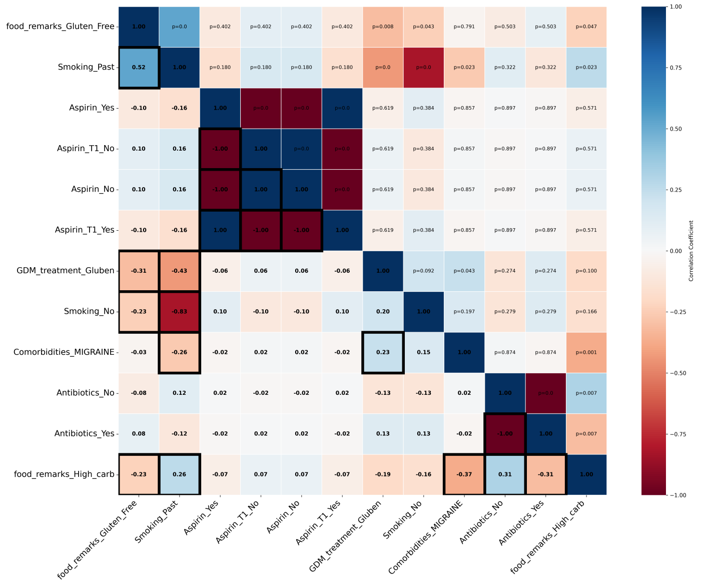


Figure S28: Correlation matrix of Israeli maternal features during the third trimester (T3). Fewer significant inter-feature correlations, reflecting reduced behavioral variability late in pregnancy.
